# *Ex vivo* autologous immune cell assay to study modulation of type 2 Inflammation by processed- or novel food extracts in peanut allergy

**DOI:** 10.64898/2026.02.13.705773

**Authors:** Robine Janssen, Alinda J. Berends, Marit Zuurveld, Severina Terlouw, Govardus A.H. de Jong, Dianne B.P.M. Somhorst, Anouk Boudewijn, Sharon Veenbergen, Harry J. Wichers, Johan Garssen, Shanna Bastiaan-Net, Rosalinde Masereeuw, Nicolette W. de Jong, Linette E.M. Willemsen

## Abstract

Peanut allergy represents a major food-allergy burden, raising concerns about food processing and novel dietary products. Current diagnostics assess primarily allergic endpoints rather than immune mechanisms initiating and maintaining type 2 inflammation, particularly DC2-mediated Th2 polarization. Here, an *ex vivo* autologous type 2 inflammatory monocyte-derived dendritic cell (moDC) DC2/T cell and B cell assay has been introduced enabling to study immunomodulatory effects induced by unprocessed (P-D) versus processed (P-DH (heated) or P-DHG (heated and glycated)) peanut proteins. Separately, different emerging food samples were assessed (protein concentrates or whole biomass) enabling to study their capacities to modify inflammatory DC2/T cell responses. CD14⁺ monocytes, CD4^+^ T cells and CD19⁺ B cells were isolated from six peanut-allergic patients’ PBMCs. MoDCs generated with IL4/GM-CSF were exposed (48h) to type 2 polarizing cytokine (DC2) mix, or DC2 mix combined with the food samples. Next, DC2s were co-cultured with T cells (5d), followed by B cells incubation with DC2/T cell supernatant and food samples (10d). Supernatants and cells were analyzed for Th1/Th2/Th-regulatory (Treg) cells, IgE and IgG profiles. DC2 induced a strong Th2 phenotype and activity, P-D DC2 further enhanced IL13 secretion and %Tregs. P-DH DC2 favored Th2, whereas P-DHG DC2 increased IFNγ, with neither increasing %Treg. P-D and P-DH DC2 increased CD40L^+^CD25^+^ memory Th2, and P-DHG showed the same tendency. Wheat, whey and seaweed biomass had little effect on DC2/Th2 driven type 2 inflammation at the dose studied, whereas algae showed distinct immunomodulatory, adjuvant-like activity upon co-exposure during inflammatory DC2 maturation. In conclusion, this autologous in vitro assay captures modification of type 2 inflammatory response in DC2-Th2 interaction, which can compile both allergen-specific and generic reactivity to food samples, supporting the potential future use of this assay as an additional tool to assess type 2-driving potential and possible allergenicity of emerging foods and processing methods in peanut-allergic patients.

**Graphical abstract:** 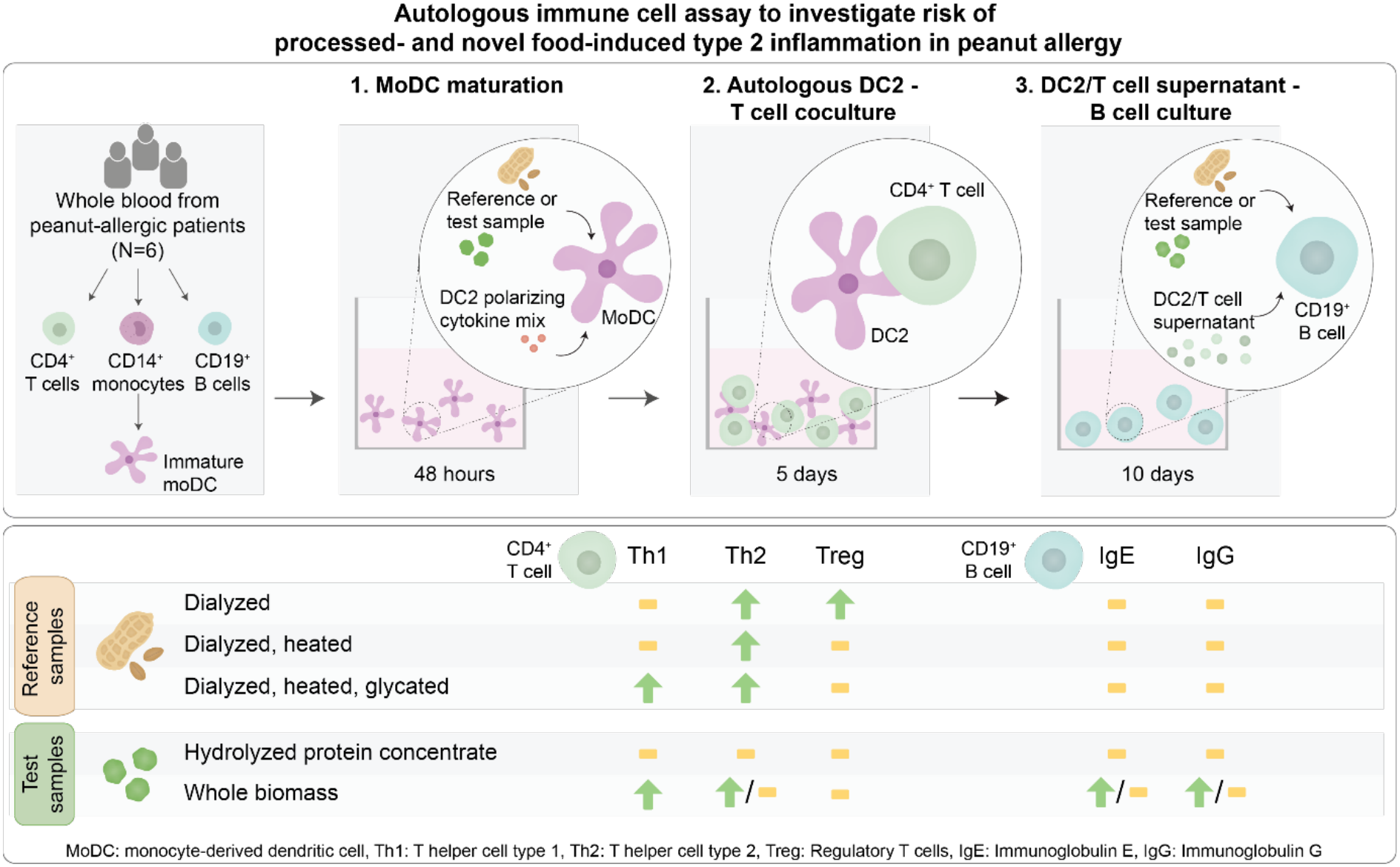

Immune cell and other illustration elements were adapted from (1, 2) and (54). Illustrations from (1, 2) were adapted with permission; permission conveyed through Copyright Clearance Center, Inc. Reference (1): © The Authors, published by Wiley-VCH GmbH under a CC BY-NC-ND license. Reference (2): © The Authors, published by Elsevier Ltd under a CC BY 4.0 license. Illustrations from (54) were adapted under the terms of the CC BY 4.0 license; © The Authors, published by Wiley-VCH GmbH.

**Capsule summary:** The autologous moDC-T cell-B cell *in vitro* assay may be used to assess whether processing methods or new foods might have intrinsic capacity to affect type 2 inflammation in peanut-allergic patients.

## 1. Introduction

Peanut allergy is one of the most common and severe food allergies, affecting 1–2% of individuals in Western countries, mostly initiated during childhood and persisting into adulthood, with significant impact on quality of life (3). Contributing factors include shifts in infant feeding, reduced and/or abnormal early-life microbial exposure, environmental pollutants, and (highly) processed food consumption (4, 5). These trends highlight the urgent need for improved diagnostics, early interventions, and tailored long-term management, as well as rigorous testing of novel proteins and biomass for safety and allergenicity as recommended by food safety authorities (6).

Food processing, thermal or non-thermal, can change protein structure and immunogenicity, raising or lowering allergenicity; however, it often does not fully remove immune reactivity, which can affect (allergic) inflammation and/or effector responses (7, 8). Food processing has varied over time and current ultra-processed foods (UPFs), which can contain high levels of advanced glycation end-products (AGEs) due to processing, may affect allergy risk (4). With only a few human *in vitro* models available, research is needed to clarify effects on allergy risk via developing novel predictive test systems. Furthermore, peanut-allergic individuals can be multi-sensitized to other allergens (9), which may predispose them to type 2 allergic inflammation as well to newly introduced proteins. Besides, the immune system (T cell or B cell receptor) may recognize structurally similar epitopes between peanut allergens and proteins from other sources, including legumes and tree nuts (cross-reactivity) (10).

Allergic sensitization begins with antigen uptake by dendritic cells (DCs). These cells present the antigen via MHC class II to naïve CD4⁺ T cells. Together with co-stimulation and allergy-driving type 2 cytokines, this promotes Th2 differentiation and migration, yielding tissue resident Th2 effector cells that secrete especially IL4 and IL13, promoting B cell activation and class switching to produce antigen-specific IgE upon allergen encounter. Subsequently, IgE binds to FcεRI receptors on mast cells and basophils, priming them for degranulation upon re-exposure (11, 12). Allergenicity tests mainly focus on skin prick tests (SPT), allergen-specific IgE levels and subsequent confirmation by oral food challenge (OFC) tests. Recently, the European Academy of Allergy and Clinical Immunology (EAACI) guidelines for the diagnosis of IgE-mediated food allergy recommended basophil cell activation tests (BAT) as well before transitioning to the OFC (13, 14). However, these assays do not directly assess the concomitant upstream immune pathway of allergen-exposed type 2 activated DCs (DC2) promoting Th2 cell activation and maintaining the allergic inflammation cascade.

In a patient specific approach, previously a peanut specific autologous DC2/T cell assay demonstrated how fructo-oligosaccharides could influence peanut exposed DC2 maturation and modulate Th2 cell responses (15). This method enables better understanding of upstream innate and adaptive immune responses that initiate and/or maintain allergic inflammation in allergic patients and potential therapies in a personalized manner. In the current manuscript, we further developed this assay aiming at monitoring modulation of type 2 inflammation in the context of peanut processing and assessing possible immune activating properties of novel food samples and biomass in food-allergic patients. More recently, an allogeneic generic model involving allergen exposed intestinal epithelial activation, monocyte-derived dendritic cell (moDC)–T cell co-cultures, followed by T cell–B cell interactions was introduced aiming to also enable measurement of B cell IgE responses (16). Given that B cells derived allergen-specific IgE is essential for allowing allergic symptom induction by mast cells, integrating them into the previously developed autologous assays would provide information about the functional outcome of DC2-Th2 interaction.

Here, the predictive value of the autologous moDC/T cell and B cell assay is explored by studying 1) the effect of peanut protein processing on maintaining type 2 inflammation, and 2) the risk of novel food samples to induce type 2 inflammation using immune cells of peanut allergic patients.

## 2. Methods

### 2.1 Study population

From the outpatient clinic of Department of Internal Medicine, Section of Allergology & Clinical Immunology, Erasmus MC, University Medical Centre Rotterdam, heparinized whole blood samples of six different peanut-allergic patients were collected. The current *in vitro* study was conducted in accordance with the Declaration of Helsinki and approved by the Medical Ethics Review Committee of the Erasmus MC (NL79534.078.21/MEC-2021-0905) and registered in the Overview of Medical Research in the Netherlands (OMON; NL-OMON51765). All participants gave written informed consent prior to the study. Sample size was calculated on IL13 as main parameter for type 2 inflammation, based on our previous study (15). With an alpha of 0.05/4= 0.0125 and power of 0.8 and calculated effect size 2.6 a sample size of at least n=5 was found sufficient to show DC2-induced type 2 inflammation in T cells (software G*Power 3.1). Six patients between 20 and 35 years of age with an allergic reaction to peanut, confirmed by a positive skin prick test (SPT), oral food challenge or both, and IgE sensitization for multiple peanut allergens assessed by the Allergy Explorer (ALEX) multiplex array (Macro Array Diagnostics, Vienna, Austria), were included in the study (**Table 1**, **Table S1**). Patients who tolerate peanut without symptoms, have negative skin-prick and challenge tests to peanut, have used antihistamines within 72 hours before testing, are unable to discontinue beta-blockers or are using prednisone at >10 mg were excluded. A dose-finding pilot study with samples from two patients (data not shown, patients not included in the current study) was performed. We observed Th2-derived IL13 secretion following DC2 maturation combined with 10 µg/mL peanut protein, consistent with a previous report (15).

**Table 1.** Demographics and ALEX test results of peanut-allergic patients.

| **SPT<sup>a</sup>** | **Oral challenge<sup>b</sup>** | **ALEX results (kU<sub>A</sub>/L)<sup>c</sup>** |  |  |  |  |
|---|---|---|---|---|---|---|
|  |  | Peanut | Milk | Egg | Pollen<sup>d</sup> | Grains |
| + | + | + | - | - | + | - |
| + | + | + | - | - | + | + |
| NP | + | + | - | - | + | + |
| NP | + | + | - | - | + | - |
| + | NP | + | - | - | + | - |
| + | NP | + | - | - | + | + |

Patients were between 20-35 years old at blood sampling, N=3 were female and the other N=3 were male. Each row represents a different donor. <sup>a</sup>Skin prick test (SPT): positive (+) or not performed (NP) because patient was not able to stop anti-histamine tablets. <sup>b</sup>Oral peanut challenge: positive (+) or not performed (NP) because of anxiety (lost to follow-up). <sup>c</sup>Allergy Explorer (ALEX) multiplex array: negative (-) <0.3 kU<sub>A</sub>/L, or positive (+) ≥0.3 kU<sub>A</sub>/L (from which all patients reacted to at least one allergen). <sup>d</sup>Pollen refers to tree-, grass-, and/or weed pollen.

### 2.2 Peanut reference samples

For a detailed description on peanut protein preparation we refer the reader to the Supporting Information.

#### Peanut protein extract purification, heating and glycation

Defatted peanut flour was obtained as described before (17), extracted in 100 mM ammonium bicarbonate (5% w/v, pH 8), centrifuged (17,000 × g, 1 h, 20 °C), filtered, dialyzed, and stored at −80 °C. For processing, extracts (dialyzed peanut protein; P-D) were either heated (130 °C, 10 min) or supplemented with glucose for glycation prior to heating. Heated non-glycated (P-DH) and glycated (P-DHG) samples were recovered, freeze-dried, and milled for analysis. (**Table 2**).

**Table 2.** Overview of peanut protein modifications and four new dietary test samples.

| **Name/abbreviation** | **Protein source** | **Modification / matrix** |
|---|---|---|
| *Reference samples* |  |  |
| P-D | Peanut flour | Dialyzed |
| P-DH | Peanut flour | Dialyzed, heated |
| P-DHG | Peanut flour | Dialyzed, heated, glycated |
| *Test samples* |  |  |
| P1 | Sea Lettuce | Seaweed whole biomass |
| P3 | *Chlorella vulgaris* | Algae whole biomass |
| P5 | Cow’s milk | Partially hydrolyzed whey protein concentrate |
| P7 | Wheat | Partially hydrolyzed wheat protein concentrate |
| Hypoallergenic samples |  |  |
| Rubisco | Spinach | No modification after isolation |
| Gelatin | Porcine skin | No modification after purchase |

#### Protein solubility, protein content and LPS contamination

Solubility was assessed by dissolving samples (1 mg/mL in DPBS), centrifuging (300 × g, 5 min, RT), and weighing tubes before/after supernatant transfer. Peanut protein content (P-D/DH/DHG) was quantified using Pierce™ BCA Protein Assay Kit (Thermo Scientific, 23227), and lipopolysaccharides (LPS) with the Pierce LAL Chromogenic Endotoxin Quantitation Kit, according to the manufacturer’s instructions (**Table S2**).

#### SDS PAGE electrophoresis and solubility analysis

Soluble and insoluble fractions of 10 µg peanut protein in Laemmli buffer were analyzed by SDS-PAGE (BioRad) under reducing (10% β-mercaptoethanol) or non-reducing conditions, stained with 0.025% Coomassie Brilliant Blue, destained, and protein bands analyzed, which confirmed the presence of the major peanut allergens Ara h 1, 2, 3, and 6 (**Figure S**1).

### 2.3 Test samples

#### Industrial partner samples

New emerging dietary ingredients (two protein concentrates and two whole biomass samples) (**Table 2**) were provided by Cargill R&D Centre Europe, Arla Foods Ingredients Group P/S, Phycom BV, and The Seaweed Company B.V., further referred to as test samples. Because of limited patient cells available, only few conditions were tested. Selection of test samples (one out of two delivered by the industrial partners) was done without interference of the industrial partner. P1 is a whole biomass seaweed (Sea Lettuce) sample. P3 is a whole biomass algae (Chlorella vulgaris) sample. P5 is a non-filtered, partially hydrolyzed product made from whey protein concentrate. The hydrolysate was produced in a process with commercially available food grade enzymes which were inactivated upon termination of hydrolysis. P7 is a partially hydrolyzed wheat gluten protein concentrate produced with commercially available food grade enzymes and dried.

#### Protein solubility, protein content and LPS contamination

Protein content of the test samples was measured via DUMAS (Flash EA 1112 N Analyzer; Thermo Fisher Scientific, Waltham, MA, USA). Methionine was used as standard to calculate protein content within the test samples, using a protein-nitrogen conversion factor of 6.25. Solubility and LPS content was determined as described above (**Table S2**).

### 2.4 Hypoallergenic samples

**Hypoallergenic samples** rubisco (spinach) and gelatin (porcine skin) (**Table 2**) were purified as described (18) or purchased (Sigma-Aldrich, G2500) respectively. Protein content was measured with the BCA assay, and LPS content as described above (**Table S2**).

### 2.5 MoDC/T cell and B cell assay

#### Immune cell isolation

PBMCs were isolated from heparinized blood in Leucosep tubes, washed, and separated into CD14⁺ monocytes, CD19⁺ B cells, and CD4⁺ T cells using magnetic-activated cell sorting (MACS). Cells were stored in liquid nitrogen or directly cultured: monocytes/moDCs in RPMI + 10% FBS + 1% pen/strep (=moDC culture medium); T and B cells in IMDM + 5% FBS, 1% pen/strep, apo-transferrin (20 µg/mL), and β-mercaptoethanol (50 µM) (=T cell culture medium). For further details we refer to the Supporting Information.

#### MoDC differentiation and maturation

CD14⁺ monocytes were differentiated into moDCs (GM-CSF, 60 ng/mL); IL4, 100 ng/mL) for six days, with medium (containing GM-CSF/IL4) refreshed every other day. On day six, one-third of the medium was replaced with stimuli as described below. MoDCs were left untreated or received a type 2 polarizing DC2 mix (prostaglandin E2 (1 μg/mL), tumor necrosis factor-*α* (50 ng/mL), IL1*β* (25 ng/mL), IL6 (10 ng/mL)) combined with either 10 µg protein/mL of peanut protein (P-D/DH/DHG), or 10 µg protein/mL hypoallergenic proteins (rubisco, gelatin), or 10 µg powder/mL of four different test samples (**Table 2**). A DC2 plus LPS (*Escherichia coli* O111:B4, Invivogen) control was included to match the highest LPS level (of rubisco; 60.85 EU/mg protein). A DC2 cytokine mixture was included because peanut-specific IL13 secretion by T cells was only detectable when moDC were maturated using this type 2 cytokine mix and not when moDC were left inactivated (15). MoDCs were incubated for 48 h (37 °C, 5% CO₂).

#### Autologous MoDC - T cell co-culture

After 48 h, matured moDCs were centrifuged (300 × g, 5 min, RT) and supernatants stored (-20 °C) for cytokine/chemokine analysis. MoDCs were resuspended in cold T cell medium, and 300,000 cells were stained for flow cytometry. The remaining 100,000 moDCs were diluted, and 60,000 were seeded in T cell medium. Autologous CD4⁺ T cells were thawed and co-cultured at a 1:10 moDC:T cell ratio (600,000 T cells). Co-cultures were incubated for five days (37 °C, 5% CO₂).

#### Intracellular T cell staining and autologous T cell supernatant - B cell culture

After five days of co-culture with moDCs, T cells were centrifuged; supernatants stored for cytokine analysis and B cell assay, and cells split for extracellular and intranuclear staining, or PMA/ionomycin (+ GolgiPlug) restimulation for intracellular cytokine staining. B cells were thawed, washed, and 100,000 cells in T cell medium were added to the T cell supernatant. Peanut reference/test samples were added to B cells. After 10 days (37 °C, 5% CO₂), with medium + food samples refreshment on day 6; supernatants (stored at −20 °C) and B cells (flow cytometry) were collected.

#### Flow cytometry – PBMC characterization

PBMCs were characterized by flow cytometry for CD4, CD45RA, CD45RO, and CCR7, with viability assessed using Fixable Viability Dye; antibody/isotype details are provided in **Table S3-4**. Before extracellular staining (30-45 min, 4 °C, dark), cells were Fc-blocked (BD Pharmingen, 564220; 10 min, 4 °C). Flow cytometry was performed on a FACS Canto II (Becton Dickinson, USA) using BD FACS Diva software (BD Biosciences) and analyzed with FlowLogic (Inivai Technologies, Australia).

#### Flow cytometry – Immune response characterization

MoDCs were analyzed for CD14, CD80, CD86, HLA-DR, CD209, OX40L, CCR7 (CD197), PD-L1 (CD274), TSLP-R (CRLF2), ST2/IL-33R, and CXCR4 (CD184), with viability assessed using LIVE/DEAD™ Fixable Near IR. 50% of the T cells were stained for CD4, CD183 (CXCR3), CD294 (CRTH2), CD69, CD25, CD40L (CD154), CXCR5 (CD185), CD196 (CCR6), CCR4 (CD194), CD161, CD45RO, and CD45RA, and intranuclear FOXP3, with viability assessed using LIVE/DEAD™ Fixable Near IR. The other 50% of the T cells were stained for CD4, and intracellular IL10, IFNγ, IL13, and IL4, with viability assessed using Fixable Viability Dye. B cells were analyzed for CD19, CD20, CD4, CD38, CD27, and CD138, and intracellular IgG and IgE, with viability assessed using LIVE/DEAD™ Fixable Near IR. Cells were Fc-blocked (10 min, 4 °C) before extracellular staining (30-45 min, 4 °C, dark), or fixed, permeabilized, and blocked before intracellular/intranuclear staining (30-45 min, 4 °C, dark). Antibody/isotype details are provided in **Table S5-12**. Flow cytometry was performed by CytoFLEX LX (Beckman Coulter, Inc. Brea, CA, USA) using CytExpert software (Beckman Coulter, Inc. Brea, CA, USA) and analyzed with FlowLogic (Inivai Technologies, Australia).

#### Enzyme-linked immunosorbent assay (ELISA)

IL10 (Invitrogen, 88-7106), IL12p70 (Invitrogen, 88-7126), CCL22 (R&D, DY366), IFNγ (Invitrogen, 88-7316), IL4 (Invitrogen, 88-7046), IL13 (Invitrogen, 88-7439), and IgE (Invitrogen, 88-50610), were measured. All ELISAs were performed per manufacturer protocols, with volumes split in halves except for blocking.

### 2.5 Statistical analysis

Data are presented as mean ± SEM and were analyzed using GraphPad Prism v10.4.1. Normality was assessed first. Paired t-tests were used for two normally distributed conditions, and RM one-way ANOVA with Dunnett’s or Tukey’s post-hoc; unequal variances were addressed by √ or log transformation or Geisser-Greenhouse correction. In limited case of missing values, mixed-effects models with Dunnett’s test was applied. Non-normal data were transformed, or Wilcoxon signed-rank or Friedman with Dunn’s post-hoc applied. Negative controls and DC2 controls were identical across matching peanut and test sample assays. Comparisons were made between the unstimulated negative control and the DC2 mix-exposed moDCs, and separately, the reference, test, and hypoallergenic samples (each combined with the DC2 mix) were individually compared to the DC2 mix alone, focusing on DC2 phenotype and DC2 function in instructing type 2 T cell responses. Given the exploratory nature of this study (N=6 different donors), these findings should be considered hypothesis-generating and warrant validation in larger cohorts. Significance was set at p ≤ 0.05 (*), p ≤ 0.01 (**), p ≤ 0.001 (***), p ≤ 0.0001 (****). When P ≥ 0.05 and ≤ 0.1, exact P-values are reported to indicate a trend.

## 3. Results

### 3.1 DC2 mix induces type 2 moDC (DC2) maturation in presence/absence of food samples

PBMCs from six peanut-allergic donors were characterized for CD4⁺ T cell subsets (naïve and memory T cells, and central and effector memory T cells) (**Figure S2**). All patients were tested peanut-positive, egg-and milk-negative, and multi-sensitized to pollen, with some reactive to seafood and grains (**Table 1, Table S1**). CD14⁺ monocytes, CD4⁺ T cells, and CD19⁺ B cells were isolated, and monocytes were differentiated into moDCs over six days. Next, moDCs were matured using a type 2 polarizing cytokine mix (DC2) alone (DC2 control) or combined with reference samples P-D, P-DH, P-DHG, test samples P1, P3, P5 or P7, hypoallergenic samples rubisco or gelatin, LPS or were left untreated (negative control) (**Figure 1A**, **Table 2**, **Table S2**). MoDC viability, absence of CD14, and dual positivity for HLA-DR (MHC-II) and CD209 (DC SIGN) (∼80% for DC2) were largely unchanged across negative control, DC2 control, and DC2 combined with reference, test, or control samples, except for P3, which reduced CD14⁻ cells by ∼25% (**Figure S3-4**). The DC2 mix increased the percentage of moDCs expressing CD80, CD86, CCR7, CXCR4, PD-L1, OX40L, and CCL22 secretion, and tended to upregulate TSLP-R, but not IL33-R (**Figure 1B-M, S3-4**). Reference P-D had no additional effect on DC2, while P-DH tended to reduce TSLP-R and CXCR4, and P-DHG decreased both plus a similar pattern for CD80, but increased CCL22 secretion (**Figure 1B, D, F-G**). CD86, CCR7, PD-L1 and OX40L remained unaffected upon exposure to P-D, P-DH, and P-DHG (**Figure 1C, E and S4 E-F**). None of the four test samples affected DC2 induced moDC maturation markers nor CCL22 secretion, except for P1 (seaweed) which slightly reduced CCL22, while P3 (algae) showed the same trend (**Figure 1H-M and S3-4**). Hypoallergenic samples generally did not affect DC2 characteristics, except for a decrease in CD86 expression with gelatin, and an increase in CCL22 secretion with rubisco but this applied also for LPS (**Figure S4**). Levels of IL10 and IL12p70 remained unchanged across reference, test, and control samples (**Figure S4Y-AD**).

**Figure 1.**
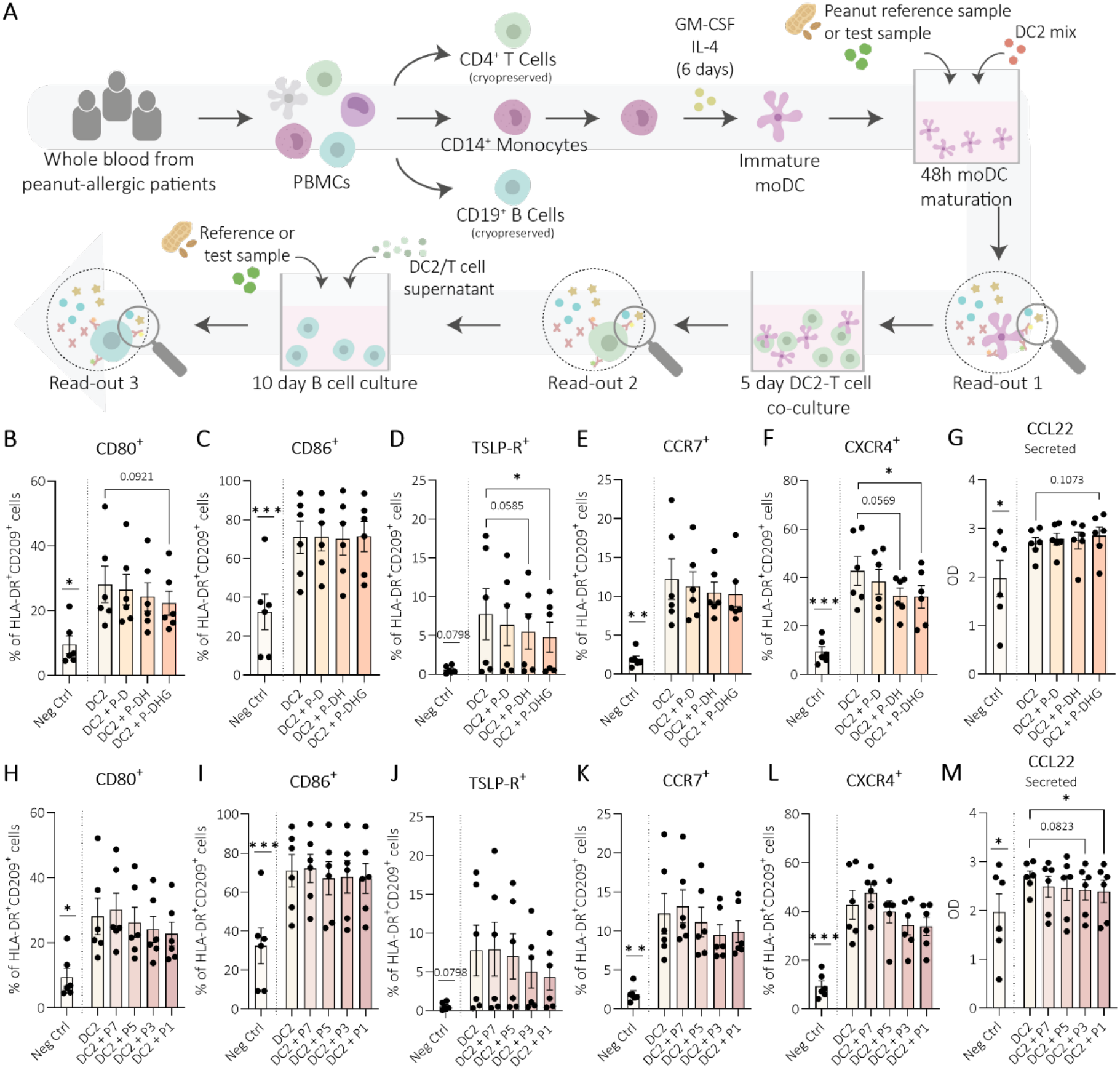
MoDCs show type 2 maturation (DC2) after DC2 cytokine mix exposure. **A**) Schematic overview of the autologous moDC–T cell–B cell assay. After 48 hours of moDC exposure to DC2 mix and **B-G**) peanut protein (P-D, P-DH, P-DHG); or **H-M**) test samples (P7, P5, P3, P1), DC2 maturation was analyzed by flow cytometry for **B, H**) CD80, **C, I**) CD86, **D, J**) TSLP-R, **E, K**) CCR7, **F, L**) CXCR4, and **G, M**) CCL22 in supernatant. Data were analyzed using paired t-test or RM one-way ANOVA with Dunnett’s test (Geisser–Greenhouse corrected where applicable). DC2 mix matured moDC (DC2) was compared to unstimulated negative control (moDC); samples (reference, test) combined with DC2 were compared to medium exposed DC2. Error bars represent mean ± SEM, N=6, p ≤ 0.05 (*), p ≤ 0.01 (**), p ≤ 0.001 (***). Immune cell and other illustration elements were adapted from (1, 2) and (54). Illustrations from (1, 2) were adapted with permission; permission conveyed through Copyright Clearance Center, Inc. Reference (1): © The Authors, published by Wiley-VCH GmbH under a CC BY-NC-ND license. Reference (2): © The Authors, published by Elsevier Ltd under a CC BY 4.0 license. Illustrations from (54) were adapted under the terms of the CC BY 4.0 license; © The Authors, published by Wiley-VCH GmbH.

### 3.2 DC2 with peanut protein or P3 shift Th2 cells from CD40L^+^CD69^+^ to CD40L^+^CD25^+^ activation

After 48 hours of moDC exposure to DC2 control or DC2 mix plus peanut reference proteins or test samples, DC2 were co-cultured with CD4⁺ T cells (**Figure 1A**). After five days, T cell viability, and CD4^+^ and CD183^-^ (CXCR3) frequency were unchanged when T cells were co-cultured with DC2 control compared to negative control, and between DC2 exposed to unprocessed and processed reference peanut protein or test samples relative to DC2 control (**Figure S5-6).** Co-culture with DC2 control increased %memory T cells (CD45RA⁻CD45RO⁺) and decreased %naïve T cells (CD45RA⁺CD45RO⁻) versus control moDCs (**Figure 2A-B, E-F**), along with elevated % CD40L (CD154) positive CD183⁻CCR6⁻CCR4⁺ Th2 cells expressing CD69 (early) or CD25 (prolonged) activation markers (**Figure 2C-D, G-H**). DC2 exposed to P-DHG further enhanced %memory Th cells on top of the effect of medium exposed DC2, while the same trend was observed for P-DH, but not for P-D (**Figure 2A**). DC2 exposed to P-D, P-DH and/or P-DHG reduced %naïve Th cells and DC2 exposed to P-D or P-DH further increased %CD40L^+^CD25^+^ memory Th2 cells compared to DC2 control, while a similar trend was shown for P-DHG (**Figure 2B, D**). P-D and P-DH DC2 tended to reduce %CD40L⁺CD69⁺ expressing Th cells compared to DC2 control (**Figure 2C, S6D**), consistent with overall upregulation of %CD40L and/or %CD25 positive CD183⁻CCR6⁻CCR4⁺ memory Th2 cells (**Figure S6E-F**). Co-culture of CD4⁺ T cells with test sample DC2 altered T cell phenotypes versus DC2 control: P3 (algae) and P7 (wheat) DC2 tended to decrease %naïve Th cells, with P3 DC2 increasing %memory Th2 cells (**Figure 2E-F**). All test samples exposed to DC2 reduced %CD40L⁺CD69⁺ memory Th2 cells (showing only a trend in P1), but only P3 DC2 increased %CD40L⁺CD25⁺ memory Th2 cells. P3 DC2 also increased %memory Th2 singularly expressing CD40L and CD25, while CD69 tended to decrease (**Figure 2G-H, S6J-L**), from which the latter was also reduced for Th cells co-cultured with P7 DC2 and P5 DC2, with a similar trend for P1 DC2. Hypoallergenic rubisco, but not gelatin nor LPS exposed DC2, similarly increased %memory, decreased %naïve T cells and enhanced %CD40L⁺CD25⁺ Th2 cells (**Figure S6**). CXCR5 and CD161 expression remained low/unaffected across samples (data not shown).

**Figure 2.**
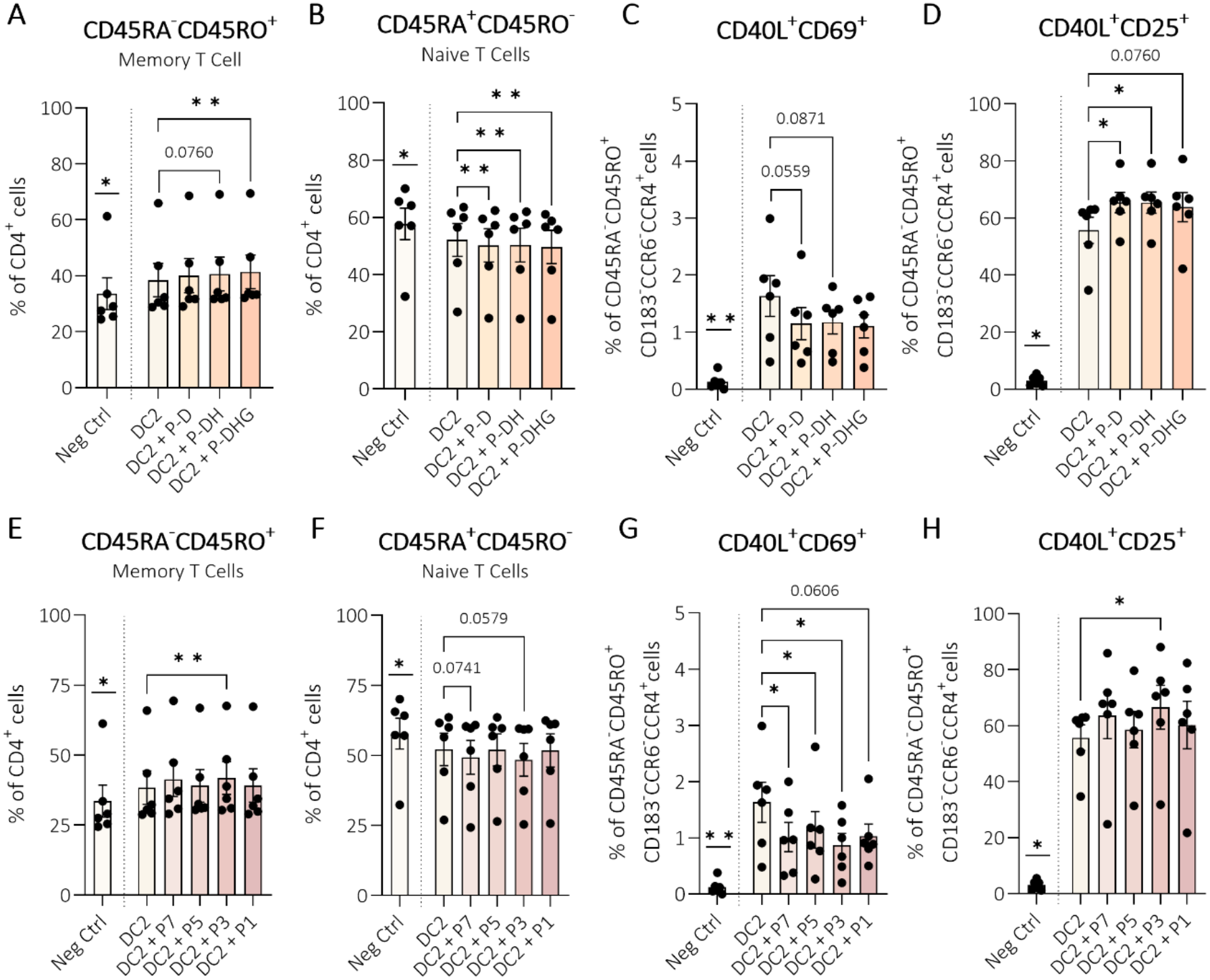
T cells co-cultured with peanut protein or specific test samples exposed DC2 shift from early to sustained activation. After 48 hours of moDC exposure to DC2 mix and **A-D**) peanut protein (D, DH, DHG) or **E-H**) test samples (P7, P5, P3, P1), DC2 were co-cultured for five days with CD4^+^ T cells. T cell phenotype was measured by flow cytometry for **A, E**) CD45RA^-^CD45RO^+^ (memory), **B, F**) CD45RA^+^CD45RO^-^ (naïve), and within memory Th2 cells: **C, G**) CD40L^+^CD69^+^, **D, H**) CD40L^+^CD25^+^. Data were analyzed using paired t-test, Wilcoxon matched-pairs signed rank test, RM one-way ANOVA with Dunnett’s test (Geisser–Greenhouse corrected), or Friedman test with Dunn’s multiple comparisons. DC2 was compared to unstimulated negative control; samples (reference, test) combined with DC2 were compared to DC2. Error bars represent mean ± SEM, N=6, p ≤ 0.05 (*), p ≤ 0.01 (**).

### 3.3 Functional Th2 responses confirmed by phenotype and IL13 secretion with peanut protein and specific test samples

Building on enhanced CD40L⁺ Th2 memory in DC2 co-culture (boosted by P-D/DH/DHG or P3 DC2), we next assessed general Th2/Th1 phenotypes and cytokines (**Figure 3, S7-S8**). Compared to co-culture with moDC negative control, DC2 control increased %CD294 (CRTH2) CD4⁺ Th cells (**Figure 3A,I**) with a similar trend for Th2 phenotype CCR6⁻CCR4⁺ in memory Th cells (**Figure 3B,J**), with higher %intracellular/secreted IL13 (**Figure 3C-D,K-L**) and %Tregs (**Figure 3H,P**). The percentage Th1 cells, based on CD183^+^ expression on CD4^+^ cells, tended to increase (**Figure 3E,M**), while secreted IFNγ decreased and intracellular IFNγ remained unaffected (**Figure 3F-G,N-O**). Compared to DC2 control, P-D DC2 left %Th2 and %intracellular IL13^+^ T cells unchanged (**Figure 3A-C**), but secreted IL13 (**Figure 3D**) and %Tregs (**Figure 3H**) increased. P-DH DC2 elevated %CD294 and %CD183^-^CCR6⁻CCR4⁺ Th2 cells (**Figure 3A-B**), with no change in intracellular IL13 (**Figure 3C**) but increased secreted IL13 by T cells (**Figure 3D**). P-DHG DC2 similarly showed an increasing pattern of %CCR6⁻CCR4⁺ Th2 (**Figure 3B**) without affecting IL13 (**Figure 3C-D**), but increased IFNγ secretion (**Figure 3G**). P-D, P-DH, or P-DHG DC2 co-cultured T cells showed unchanged %CD183 and intracellular IFNγ versus DC2 control (**Figure 3E-F**). None of the test sample DC2s affected Th2 or Th1 phenotype or cytokines compared to DC2 control, except for P3 (algae) DC2, and to a lesser extent P1 (seaweed) and P7 (wheat) DC2 (**Figure 3A-H**). Only P3 (algae) DC2 increased %CD294 (**Figure 3I**) with a similar pattern for %CCR6⁻CCR4⁺ memory Th2 cells (**Figure 3J**), without affecting %intracellular IL13^+^ Th cells (**Figure 3K**) but elevating secreted IL13 (**Figure 3L**) and both %intracellular IFNγ^+^ Th cells and secreted IFNγ (**Figure 3N-O**). P7 (wheat) DC2 tended to increase secreted IL13 (**Figure 3L**), and P1 (seaweed) DC2 increased secreted IFNγ by T cells (**Figure 3O**). LPS DC2 had little effect, while hypoallergenic rubisco-exposed DC2-T cells showed increased %Tregs, %CCR6⁻CCR4⁺ memory Th2 (**Figure S6**), and IL13 secretion (**Figure S8**), whereas gelatin DC2 increased %CD183⁺ Th1 cells compared to DC2 control (**Figure S6O**). IL10 and IL4 were low/undetectable (data not shown).

**Figure 3.**
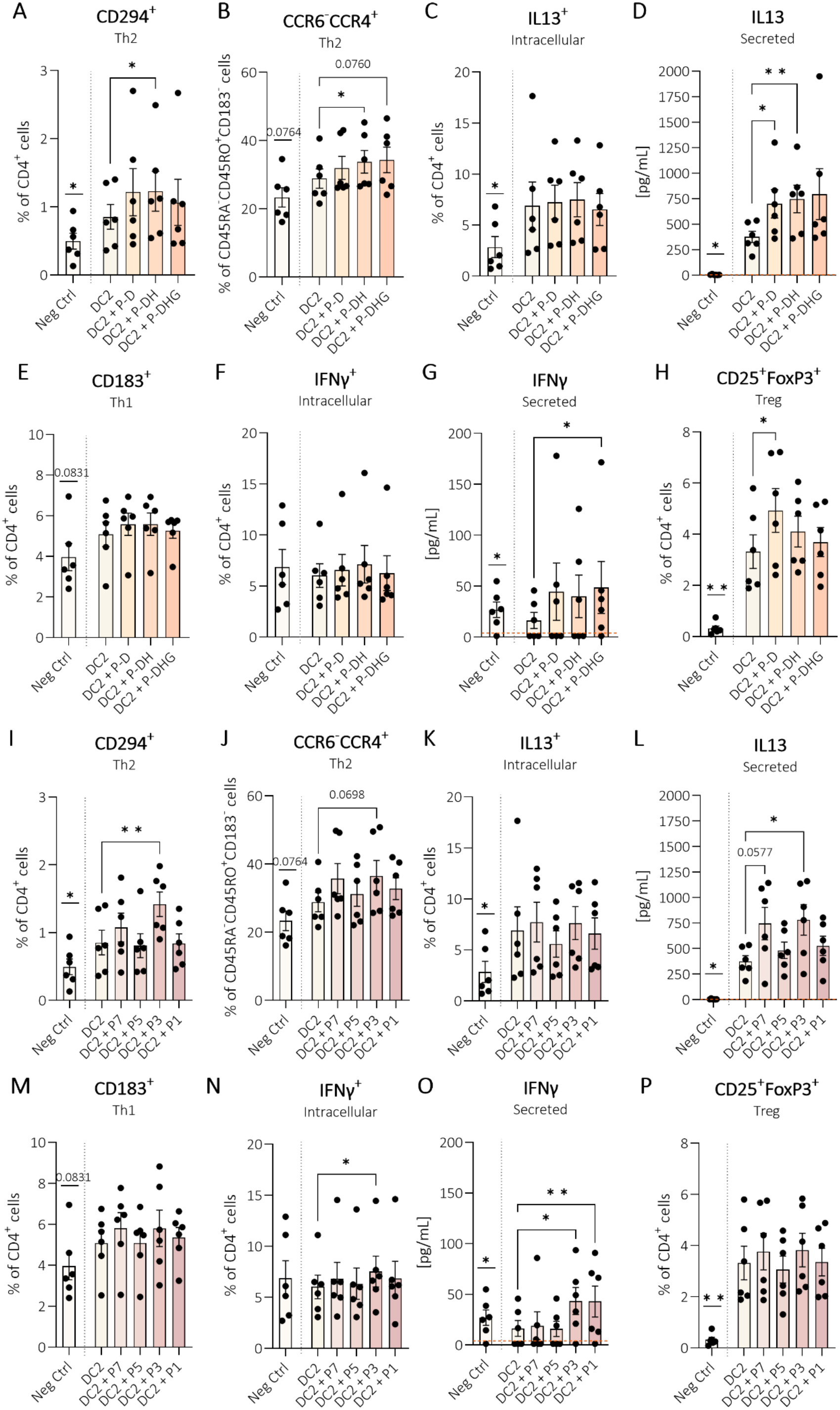
Th2 responses (phenotype and IL13) confirmed with peanut protein and selected test samples. After 48 hours of moDC exposure to DC2 mix plus **A-H**) peanut protein (D, DH, DHG) or **I-P**) test samples (P7, P5, P3, or P1), matured DC2 were co-cultured for five days with CD4^+^ T cells. Th2 phenotype was determined by **A, I**) CD294, **B, J**) CCR6^-^CCR4^+^, **C, K**) intracellular IL13, and **D, L**) secreted IL13. Th1 phenotype determined was determined by **E, M**) CD183, **F, N**) intracellular IFNγ, and **G, O**) secreted IFNγ, and Treg by **H, P**) CD25^+^FoxP3^+^. Data were analyzed using paired t-test, Wilcoxon matched-pairs signed rank test, RM one-way ANOVA with Dunnett’s test (with Geisser–Greenhouse correction where applicable), or Friedman test with Dunn’s multiple comparisons. DC2 was compared to unstimulated negative control; Samples (reference, test) combined with DC2 were compared to DC2. Error bars represent mean ± SEM, N=6, p ≤ 0.05 (*), p ≤ 0.01 (**). Orange dotted lines represent ELISA minimal detection limit.

### 3.4 Memory B cells and IgE unaffected by peanut or test samples, except algae biomass

Following 5-day DC2-T cell co-culturing, supernatants with again added peanut protein reference samples or test samples were applied to CD19⁺ B cells for 10 days (**Figure 1A**). B cells remained viable and were CD4⁻ (**Figure S9-S10**). Combined CD19⁺/CD20⁺ subsets tended to increase with supernatant of DC2-T cells compared to control moDC-T cells (**Figure S10G-**I). DC2-T cell supernatant tended to reduce %CD27 but left %CD27⁺CD38⁺ and IgG (MFI) unchanged, while IgE (MFI) tended to decrease (**Figure 4A-H**). Peanut protein DC2-T cell supernatants left B cell CD19⁺/CD20⁺ subsets, CD27⁺, CD27⁺CD38⁺ and IgG (MFI) unaffected, however, IgE (MFI) tended to decline with P-D and P-DH (**Figure 4A–D, S10G**). %CD19⁺/CD20⁺ B cells tended to increase with P3 DC2-T cell supernatants compared to control DC2-T cell supernatant, while no changes were observed for P1, P5, or P7 (**Figure S10H**). CD27 expression increased with P3 DC2-T cell supernatants (**Figure 4E**), whereas %CD27⁺CD38⁺ cells decreased (**Figure 4F**). MFI of intracellular IgE and IgG increased with P3 DC2-T cell supernatant, but not with other test samples (**Figure 4G+H**). Three donors tested positive for sensitization to grain allergens based on the ALEX multiplex array. To further indicate their responses to wheat (P7), these donors are highlighted separately (marked G+) and individual donor responses for Th2/Th1 and B cells are shown (**Figure S11**). Because most intracellular IgE and IgG percentages fell below the detection limit, MFI is shown here. However, when gating IgE versus IgG, both P3 and P1 were found to increase the % B cells expressing IgE. This effect was more pronounced for P3 (around 85% of all B cells) than P1 (around 4% of all B cells, **Figure S13C**), and only P3 increased the % B cell dual positive for IgE and %IgG (**Figure S12-13**). CD138 expression, and IgE secretion, were for all samples low or not detected (data not shown). Hypoallergenic and LPS controls did not affect any of the markers, except for rubisco, which showed an increasing trend for IgG MFI (**Figure S10I-M**).

**Figure 4.**
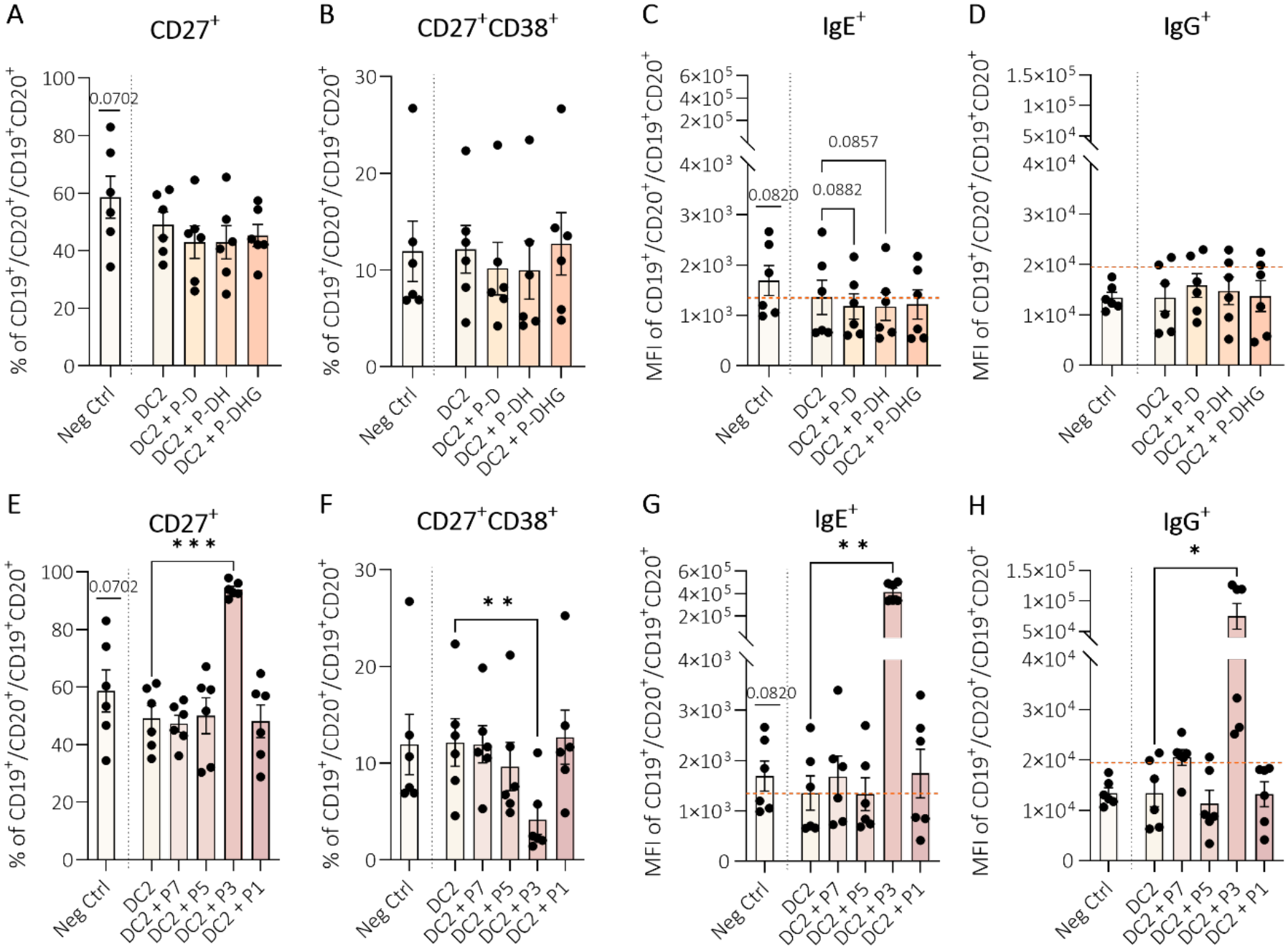
Memory B cell and IgE were unchanged by peanut DC2-T cell supernatants but increased by algae biomass. After five days of control, reference or test sample exposed DC2–T cell co-culture, CD19^+^ B cells were incubated with the co-culture supernatant with **A-D**) extra added peanut protein (P-D, P-DH, P-DHG) or **E-H**) test samples (P7, P5, P3, P1) for 10 days. B cells were analyzed for **A, E**) %CD27^+^, **B, F**) %CD27^+^CD38^+^, and MFIs of **C, G**) IgE, **D, H**) IgG. Orange dotted lines represents average MFI of corresponding FMO. Data were analyzed using paired t-test, RM one-way ANOVA with Dunnett’s test (with Geisser–Greenhouse correction where applicable), or Friedman test with Dunn’s multiple comparisons. DC2 was compared to unstimulated negative control; Samples (reference, test) combined with DC2 were compared to DC2. Error bars represent mean ± SEM, N=6, p ≤ 0.05 (*), p ≤ 0.01 (**), p ≤ 0.001 (***).

## 4. Discussion

This study used an autologous DC2-T cell *in vitro* assay for testing type 2 driving capacity of processed peanut protein (P-DH and P-DHG) as well as novel industrial proteins and whole biomass (test samples). In addition, autologous B cell responses to DC2-T cell supernatants in the presence of these food samples were examined. MoDC exposed to DC2 mix induced clear Th2-polarizing activity, which was further enhanced in case also naïve peanut protein (P-D) was added to the DC2 mix, yielding increased IL13 secretion and Treg frequency. P-D processing modified the type 2 directed T cell response. Heat treatment (P-DH) further favored the Th2 phenotype, while losing the peanut induced increase in %Treg. In contrast, even though peanut heat treatment and glycation (P-DHG) also resulted in loss of Treg induction, it enhanced type 1 IFNγ release in association with reduced co-stimulatory marker expression on DC2. DC2 exposed P-D, P-DH, with a similar pattern for P-DHG, increased late activated memory Th2 cells compared to DC2 control. Separately, diverse test samples were studied for their immunomodulatory effect. Among these, P7 (wheat), P5 (whey), and P1 (seaweed) showed no substantial effects in the dose studied, whereas P3 (algae) exposed DC2 displayed distinct immunomodulatory activity promoting both type 2 and type 1 T-and B cell responses, suggestive of a more general adjuvant-like function.

Alongside concomittant exposure to peanut reference and test samples, type 2-maturing DC2 mix was compared to unstimulated moDC controls first. Consistent with *Hayen et al.* (15), it was observed that DC2 mix similarly upregulated DC2 CD80 and CD86 expression, but did not induce IL10 secretion. Beyond these markers, DC2 also increasingly expressed type 2 related TSLP-R, while secreting CCL22. Furthermore, migration markers CCR7 and CXCR4 increased, known to be required for trafficking to the lymph nodes to initiate antigen specific adaptive immune responses. (16, 19, 20) DC2 induced IL13 secretion and promoted CD294 (CRTH2) expression in T cells, while CD183 (CXCR3) remained unchanged and IFNγ secretion was reduced, confirming both Th2-polarizing activity and assay reliability. These Th2-polarizing effects in absence of peanut protein are likely generic, known to occur also in DC2-T cell co-cultures using non-allergic donors.

Combined exposure of moDC to DC2 cytokines mix and peanut protein (P-D DC2) did not affect DC2 phenotype nor enhanced Th2 phenotypic markers, but further increased IL13 secretion by T cells compared to control DC2, this may be indicative of a peanut specific response, consistent with earlier findings. (15) In addition, early activation (CD40L⁺CD69⁺) of memory Th2 cells tended to decrease, whereas the late activation marker (CD40L⁺CD25⁺) increased, possibly indicative of responsivity of allergen-specific activated Th2 cells of peanut-allergic patients (21, 22) upon P-D epitope presentation via DC2. The latter was the case for processed P-D also, indicating that T cell epitope recognition was conserved during processing. Beyond increasing IL13 secretion, P-D DC2 enhanced Treg (CD25⁺Foxp3⁺) frequency also. However, we cannot exclude a generic effect of the peanut protein extract as also other exposures, including assumed hypoallergenic rubisco and biomass sample P3, enhanced IL13 secretion on top of the effect of medium-exposed DC2 on T cells. It would be of interest to compare sensitivity of cells from non-allergic donors with those of peanut allergic patients in future studies, to show that indeed peanut exposed DC2 as assumed only provoke a Th2 response in cells of allergic patients having TCR recognition for peanut protein epitopes. However, it is also of interest to compare responsiveness of cells of peanut allergic patients with cells of non-allergic individuals in case of novel food samples’ responsiveness as cells of these patients may be more sensitive to either generic or allergen-specific effects.

Heating or heating and glycation of peanut proteins modulated DC2-T cell responses. Even though both prevented a rise in %Treg, P-DH maintained the type 2 signature while P-DHG promoted type 1 IFNγ secretion. P-DH minimally impacted DC2-T cell activation, suggesting that heating alone does not strongly alter maturation of DC2, yet it promoted Th2-phenotypic marker expression while maintaining enhanced Th2 activity (IL13), consistent with late memory Th2 cell activation (CD40L⁺CD25⁺). This indicates potential prolonged immune engagement. P-DHG DC2 did affect DC2 phenotype, lowering %CD80, TSLP-R and CXCR4 expressing DC2, while tending to increase CCL22. Hence, P-DHG modified DC2 co-stimulatory and chemokine signals, which could lead to differential T cell priming (16, 19, 20). The functional outcome of the P-DHG DC2-T cell response was the promotion of Th1 (IFNγ) response. Mechanistically, these P-DH and P-DHG induced immune alterations via affecting DC2 function, may be linked to structural changes of allergenic proteins induced by processing, as e.g. heating denatures and aggregates Ara h 2 and Ara h 6 (23). Because heating often lowers IgE binding e.g. by disrupting conformational B cell epitopes, denatured allergens often show reduced allergenicity as assessed by effector cell assays such as BAT (7). However, the autologous DC2-T cell assay allows to study the effect of allergen processing in upstream processes maintaining type 2 inflammation, either via altered generic modulation of DC2 function or via altered processing and presentation of linear T cell epitopes, hereby modifying T cell responses. Roasting at high temperatures is known to enhance allergenicity, likely through the formation of AGEs (7). Importantly, these AGEs, formed during the Maillard reaction widely applied in food industry, result in protein modifications that can also modify function and phenotype of antigen-presenting cells (24). This provides a plausible explanation for the altered P-DHG DC2 activation, shaping the DC2 phenotype and consequent T cell priming. The DC2-T cell assay shows that, eventhough heating and glycation of peanut protein promotes type 1 which may couteract type 2 inflammation, Treg promoting activity is lost, which is unwanted since this means loss of tolerance inducing capacity. These counterbalancing responses to peanut exposure have been observed before, e.g. with Omalizumab skewing Th1 upon peanut oral immunotherapy but then without Treg changes (25). Alternatively naïve peanut protein not only induced type 2 but also Treg frequency. Moreover, a delayed and persistent Treg activation has been observed in human upon OFC (26). However, the molecular changes that underlie these altered APC-T cell responses upon P-DH and P-DHG exposure remain to be elucidated in future studies.

Novel food samples showed no DC marker changes, though P1 (seaweed) significantly reduced CCL22, while P3 (algae) showed the same tendency. In the functional DC2-T cell outcome, P1 DC2 were shown to selectively promote type 1 IFNγ release by T cells, while P3 DC2 promoted both IL13 and IFNγ secretion. P3 also shifted the naïve/memory T cell and CD69^+^/CD25^+^ CD40L⁺ memory Th2 subsets, and increased both B cell-derived IgE and IgG, indicating strong immunomodulatory effects. Other test samples reduced CD40L⁺CD69⁺ without sustained memory Th2 cell shifts towards a CD40L^+^CD25^+^ phenotype, suggesting limited Th2 activation with wheat (P7), whey (P5) or seaweed (P1). Yet, P7 (wheat)-DC2 did tend (not significantly) to promote IL13 secretion by T cells. Wheat protein may have modified DC2 function to support Th2 activation or may contain allergens that crossreact with T cells of peanut allergic patients. Peanut (legume) proteins (e.g., Ara h 2, legumins, vicilins) are known to cross-react with soy, hazelnut and birch pollen, but cross-reactivity with wheat, whey, seaweed or algae is expected to be low or absent (27, 28). However, peanut allergic patients are often multisensitized and all were also pollen sensitized. Cross-reactivity between pollen (grass) and wheat (grain/grass) is described (29, 30). Wheat epitopes presented by DC2 may therefore have been able to activate pollen-specific T cells, tending to support type 2 inflammation. Alternatively, there may be a generic effect of undigested wheat protein concentrate on moDCs of food-allergic patients enhancing DC2-induced T cell activation. In this study the test samples were added on weight base, meaning that protein content within each food matrix differed considerable among samples. In addition, protein solubility differed between samples **(Table S2)**, therefore the absence of a response to P1, P5, and P7 cannot be interpreted as low biological activity of the protein content. Further dose-finding studies are required to assess this.

P3 DC2 promoted a Th2 signature (increased %CD294⁺ and %CCR6^-^CCR4^+^ T cells, and IL13 secretion), and also enhanced type 1 IFNγ (intracellular and secreted) in T cells, implying a balanced immune profile. Interestingly, P1 DC2 selectively promoted type 1 IFNγ secretion by T cells. In these cases, overall immune responses likely arose from whole biomass rather than isolated proteins, pointing to possible adjuvant effects of cell wall components or lipids. Microbial-or pathogen-associated molecular patterns present on algae, roots, and herbs can activate pattern recognition receptors (PRRs) and boost immunity (31), an effect that likely was observed for P3 and P1. P3 DC2 promoted both type 2 and 1 responses and similar adjuvant activity is known in pollen allergy, where lipids, carbohydrates and microbiota can enhance immune activation via PRRs, including C-type lectin receptors (32). Supporting this, marine algae extracts at comparable concentrations (0.1-20 µg/mL) have been shown to induce pro-inflammatory antigen presenting cell profiles, T cell proliferation and IFNγ production (33). Thus, immune effects of whole algae biomass may reflect generic adjuvant-like properties. While algae-and seaweed-related allergies are rare, the broad immune-stimulating effects of P3 warrant further study as a potential adjuvant or immune booster, with nutritional benefits carefully balanced against the risks. However, it should be considered that this possible “adjuvant-like” effect was observed when the sample was combined with the DC2-mix (inflammatory cytokines), which already activates immune cells toward type 2 inflammation. In future studies, these immune stimulating effects by algae and seaweed biomass should be studied further using also cells from non-allergic individuals, and under both control as well as type 2 inflammatory conditions.

Besides the immune-stimulating effects described above, B cells were exposed to DC2-T cell supernatant plus P3 were activated to express IgE and IgG, while naïve or heated peanut protein extract or P5 or P7 left B cells largely unaffected. P1 did not significantly enhance MFI, but a small percentage of B cells were found to express IgE, when analyzing IgE and IgG expression (**Figure S13 C+G**). For P3, the elevated MFI was also reflected by the proportion of IgE-positive B cells, as well as in the percentage of B cells double positive for IgE and IgG. This co-expression may reflect cells caught mid-switch, transitioning from IgG to IgE production (34, 35). Alternatively, it might stem, at least in part, from secreted IgE and IgG antibodies attaching to the B-cell surface. As the intention was to measure intracellular immunoglobulin expression, such membrane-bound and receptor-bound antibodies would first need to be enzymatically cleaved off (e.g., via trypsin treatment) before fixation and permeabilization; without this step, surface-bound IgE and IgG may have inadvertently been captured alongside the intracellular signal, contributing to the observed double positivity (36, 37).

As P3, and to a more limited extent P1, unexpectedly raised the frequency of IgE-positive B cells, and P3 also increased the IgE/IgG double-positive population, this is notable given that B cell activation typically requires BCR-allergen engagement combined with T cell contact (MHC-II presentation, CD40/CD40L co-stimulation, T cell-derived cytokines), which can be further enhance by TLR signaling (38, 39). Since our assay involved only DC2/T cell supernatant plus proteins or biomass, without direct T cell contact or anti-CD40, the IgE response observed must have arisen independently of T cell contact. Memory B cells can indeed be activated this way, mainly through TLR engagement (40), or through strong BCR cross-linking by repetitive structures like polysaccharides combined with TLR or cytokine signaling (41, 42). Future work could add isolated memory T cells to memory B cells from these cultures to further dissect protein-specific effects. Additionally, further characterization of algae and seaweed in future work could help clarify whether autofluorescence or other optical properties might have influenced the measured signal. Although this potential confounding effect was likely limited by our standard gating strategy, including gating on cells and singlets as well as cell-specific markers and no signal was observed in the FITC channel which can detect autofluorescence. Moreover, in the current study, allergen-specific IgE was not measured and needs further clarification in future studies.

*In vivo*, only bypassing the intestinal barrier would allow proteins to directly interact with immune cells (12, 19), which we mimicked in this study without prior digestion. Normally, gastrointestinal digestion breaks down proteins into peptides and amino acids for absorption, while processing (e.g., heating and/or glycation) can alter their structure and bioavailability (43). Instead, we recreated a type 2-driving environment by culturing moDC with a DC2 cytokine mix, though translation to the *in vivo* setting remains uncertain. Incorporating an epithelial barrier could strengthen future models (2). In our model, we did not only address effects of whey and wheat protein concentrates, we also introduced whole biomass seaweed and algae. The protein content of these samples was limited, therefore, adjuvant effects may be linked to other matrix molecules. To produce their direct adjuvant effects, crossing the barrier would be a requisite. However, algae remain structurally intact during processing and are relatively large in size (e.g., green microalgae have a size of ∼1-10 µm (44)). Tight junctions in the intestinal barrier normally restrict passage to small molecules (∼0.6 nm diameter) (45), whereas compromised tight junctions allow for protein passage up to ∼12.5 nm in diameter (19, 45). Accordingly, paracellular transport of intact algae across epithelial barriers is highly unlikely due to their dimensions. This may explain why eventhough in the current model P3 DC2 greatly promoted type 1 and 2 responses in both T-and B cells, algae food allergies are relatively uncommon.

We included rubisco as a putative hypoallergenic protein. However, rubisco DC2, in addition to increasing Tregs frequency, also promoted the Th2 phenotype and IL13 secretion, suggesting that rubisco may not be fully non-allergenic as anticipated. Rubisco is a plant-derived protein present in green leafy vegetables and is generally considered to have a low allergenic potential (46). However, industrial isolation of rubisco can be technically challenging (47), and purified rubisco is, therefore, not yet widely available in food products. In our assay, however, rubisco promoted type 2 inflammation, suggesting that this protein may not be fully immunologically inert. Given that rubisco is known for its high digestibility, in future studies it would be of interest to characterize its type 2-driving properties by 1) comparing whole versus digested protein, 2) comparing different industrial isolation procedures, and 3) testing rubisco in non-activated cells from non-allergic individuals to determine whether peanut-allergic patients are more responsive to isolated rubisco than non-allergic controls. Furthermore, green algae are reported to contain rubisco (48), which might explain the immune effects observed in test sample P3. The rubisco sample contained more LPS compared to other samples, although the amount was still very low (∼61 pg/mL, **Table S2**). In addition, the DC2+LPS control, matched for the highest LPS level, did not reproduce rubisco’s effects on memory Th2 cells, CD40L⁺CD25⁺ cells, or IgG. These findings indicate that the observed immune activating effect by rubisco is unlikely to be explained by LPS contamination alone. The component within the rubisco sample responsible for this effect was not further resolved in the present study. Thus, rubisco-related immune effects point to an intriguing avenue for future research on rubisco-containing foods under allergic conditions, as modeled in our *in vitro* assay. In addition, a remaining question, particularly for the food test samples, is whether all proteins were immune-accessible, as some may be trapped in the matrix but can be released upon digestion. Furthermore, we used heat-inactivated FBS as the serum type for all conditions to avoid introducing an additional source of variability between patient samples, and because previous studies showed comparable results when using heat-inactivated FBS or pooled human serum in similar assays using human immune cells (49, 50). Also similar to our previous studies using X-VIVO^®^ 15 medium enabling serum free culturing (15), we show a comparable effect of peanut exposed DC2 driving Th2 activation using heat-inactivated FBS. However, a contribution of FBS to the observed effect cannot be fully excluded. In future studies, this could be optimized by using patient-derived pooled or specific serum to improve the translational value of the observations. However, this would introduce additional variability between patient samples, warranting careful validation and characterization of the serum, as well as careful analysis and interpretation of the results across patients. Although peanut (processed or unprocessed) had little effect on B cell-derived IgE or IgG responses, IgE was not increased with unprocessed and heated peanut. The lack of robust IgE production, despite Th2 responses, may reflect the low frequency of allergen-specific B cells (∼0.01% Ara h 2/6-binding (51)) in our mixed naïve/memory CD19^+^ pool and the use of only the food samples and supernatant from activated T cells, with T cells excluded to prevent naïve B cell activation. Future work focusing on direct T-memory and B cell interactions may help clarifying these pathways and potentially amplifying B cell responses.

To summarize, this autologous *in vitro* assay demonstrates peanut exposed inflammatory DC2 to enhance Th2 responses in cells of peanut allergic patients, and its modulation by peanut protein processing. In addition, differential reactivity to food test samples was observed in this DC2-Th2 model, enabling studies on intrinsic capacities of protein concentrate or biomass to enhance or modify inflammation. This assay was based on a limited sample size of six peanut-allergic patients and is, therefore, explorative and hypothesis-generating in nature, with results providing an indication of possible type 2 inflammation modulation. Future validation should be performed in larger peanut-allergic patient cohorts, and the assay should be extended to other food-allergic groups (e.g., hen’s egg-or cow’s milk-allergic patients) along with non-allergic controls. This will help establish the utility of this autologous assay as a potential complement to existing clinical tests that monitor end-stage effector responses (SPT, IgE, OFC, BAT), by assessing upstream processes of DC instruction of Th2/Th1/Treg balance and its consequences for type 2 inflammation. Furthermore, once further validated and extended to common food allergies such as hen’s egg or cow’s milk, this platform may provide a foundation for testing additional food-allergic patient groups and evaluating other food processing methods and novel food protein sources or biomass. In addition, generic, *de novo* and antigen-specific responses should be separated in such an assay. In addition, further refinement may be introduced by only including naïve or memory T cells (and B cells) depending on the research question, rather than the combination as used in our assay. An *ex vivo* assay may also allow testing of a larger number of samples than is feasible *in vivo*, with lower burden for patients compared to OFC or SPT, allowing to evaluate novel foods or processing methods that have not yet been recognized as safe (GRAS) (52). With potential applications in personalized food allergy assessment and broader food-safety evaluation (including groups at-risk such as those with atopic dermatitis (53)), this platform could be validated further to potentially complement current allergenicity approaches, which predominantly focus on upstream effector DC2/Th2-cell responses relevant in initiating and maintaining allergic inflammation. Furthermore, it may contribute to clinical diagnostics and to a more mechanistic understanding of immunomodulatory properties of processed and novel food proteins.

## Acknowledgements

We would like to thank the knowledge partners in the consortium (Wageningen UR dep. Wageningen Food & Biobased Research, Erasmus MC dep. Allergology and clinical immunology, and Utrecht University div. Pharmacology) for their substantive scientific contribution and for having made this project possible. We also acknowledge the Dutch funding program: This project receives financial support from the Top Sector Agri & Food. Within the Top Sector, the business community, knowledge institutions and the government work together on innovations for safe and healthy food for 9 billion people in a resilient world. We also would like to acknowledge the industrial partners Danone Nutricia Research B.V., Cargill R&D Centre Europe, Arla Foods Ingredients Group P/S, Société des Produits Nestlé S.A., Phycom BV, The Seaweed Company B.V., and Mycorena AB. The authors would like to thank S. Thijssen (Department of Pharmaceutical Sciences, div. Pharmacology, The Netherlands) for the support in Flow Cytometry analysis, J. Visser for the support in conducting ELISAs (Department of Pharmaceutical Sciences, div. Pharmacology, The Netherlands), M.M.M. Tomassen (Wageningen UR dep. Wageningen Food & Biobased Research, The Netherlands) for the contribution to LPS measurements, and T. Hoppenbrouwers (Wageningen UR dep. Wageningen Food & Biobased Research, The Netherlands) for the support in patient study design.

This version was revised in response to reviewers’ comments received during peer review at *Frontiers in Immunology*.

## Author Contributions Statement

R.J. and L.E.M.W. designed the *in vitro* study. R.J. and A.J.B. performed the *in vitro* experiments. M.Z. supported in the flow cytometry experiments. R.J. analyzed the data. M.Z. supported in flow cytometry data analysis. N.W.J. and L.E.M.W. set up the patient study design and wrote the METC. N.W.J. included all peanut allergic patients and N.W.J and S.T. obtained blood samples. M.Z., S.B-N, and R.M. supported in *in vitro* study design. G.A.H.J. isolated and provided peanut protein and hypoallergenic rubisco. D.B.P.M.S contributed to DUMAS measurements and protein solubility measurements. D.B.P.M.S and A.B. contributed to heating and glycation of the peanut protein. S.V. contributed in collecting patient information and test results. H.J.W. and J.G. contributed in securing funding. R.J. wrote the manuscript. S.B-N, R.M. and L.E.M.W. reviewed and edited the manuscript. All authors read and contributed to the final version of the manuscript.

## Data availability statement

The raw data supporting the conclusions of this article will be made available by the authors upon reasonable request.

## Abbreviations

AGEs: Advanced glycation end-products
ALEX: Allergy Explorer (ALEX) multiplex array
BAT: Basophil activation test
CCR: C-C chemokine receptor
CD: Cluster of differentiation
CXCR: C-X-C chemokine receptor
DCs: Dendritic cells
ELISA: Enzyme-linked immunosorbent assay
HLA-DR: Human leukocyte antigen – DR
IFNγ: Interferon gamma
IgE/IgG: Immunoglobulin E / Immunoglobulin G
IL: Interleukin
LPS: Lipopolysaccharides
MFI: Median fluorescence intensity
FMO: Fluorescence Minus One
METC: Medical Ethics Review Committee
MHC-II: Major histocompatibility complex class II
MoDC: Monocyte-derived dendritic cell
NP: Not performed
OFC: Oral food challenge
OX40L: OX40 ligand
PBMC: Peripheral Blood Mononuclear
Cells P-D: Dialysed peanut protein
P-DH: Dialysed and heated peanut protein
P-DHG: Dialysed, heated and glycated peanut protein
PPR: Pattern recognition receptor
P1: Sea Lettuce (Seaweed whole biomass)
P3: Chlorella vulgaris (Algae whole biomass)
P5: Cow’s milk (Hydrolyzed whey protein concentrate)
P7: Wheat (Hydrolyzed Wheat protein concentrate)
PD-L1: Programmed death-ligand 1
SEM: Standard error of the mean
SPT: Skin prick test
Th1: T helper type 1 (cell)
Th2: T helper type 2 (cell)
UPFs: Ultra-processed foods

