## Supporting Information for "*Ex vivo* autologous immune cell assay to study modulation of type 2 Inflammation by processed- or novel food extracts in peanut allergy"

**Detailed Experimental Section 1: Peanut reference sample proteins**

**Peanut protein extract purification**

Defatted peanut flour, from cultivar runner market-type, was obtained according to the method described before (1). The peanut flour was dispersed into 100 mM ammonium bicarbonate (5% w/v, pH 8), stirred at room temperature (RT) for 1 hour, and centrifuged at 17,000 × g for 1 hour at 20°C. The obtained supernatant was filtered Whatman Schleicher & Schuell filter paper (595½). The filtered supernatant was dialysed against 100 mM ammonium bicarbonate buffer (pH 8) at RT (675 mL protein extract : 20 L dialysis buffer) using 3.5 kD MWCO dialysis tubes (Spectra/Por™ 3 Standard RC tubing; Fisher Scientific, Landsmeer, The Netherlands). After 17 hours, dialysis buffer was refreshed, and dialysis was continued for another 5 hours. The dialysed supernatant was again centrifuged at 17,000 × g for 1 hour at 20°C, and the resulting supernatant was snap frozen in liquid nitrogen and stored at -80°C. One part was freeze dried resulting in sample P-D, the remaining supernatant was used to create the processed peanut extracts (**Table 2**).

**Heating and glycation of peanut protein**

Peanut protein extract was heated under dry conditions, with or without addition of glucose monohydrate (Sigma-Aldrich Chemicals, The Netherlands). Per gram of protein, an equal molar amount of glucose was added per lysine content to ensure proper glycation of protein samples. Peanut protein contains 39.2 mg or 0.31 mmol of lysine per gram of protein (2). Samples were prepared as follows: dialyzed peanut samples were thawed at 37°C to guarantee solubility of all allergens. To half of the samples, glucose monohydrate was added in the ratio of 55.8 mg glucose per gram of protein. All samples were then transferred to screw-cap glass test tubes (Schott GL18) and freeze dried using an Edwards Lyofast S 08 Freeze Dryer (A. de Jong TH, Papendrecht, The Netherlands). Afterwards, the samples were stored in a glass desiccator filled with saturated sodium bromide solution for two weeks at 20°C in order to obtain water activity (aw) of 0.59 (3). All samples were then heated at 130°C for 10 minutes in a Reacti-Therm III #TS-18823 heating module (Thermo Fisher Scientific Inc, The Netherlands). After heating, the samples were immediately placed in an ice bath to cool down immediately. After cooling, proteins were reconstituted in 100mM ammonium bicarbonate (pH 8). The peanut protein samples were stirred for 1 hour at room temperature. Afterwards, heated but non-glycated samples (DH-samples) were snap frozen in liquid nitrogen and freeze dried. The glycated samples (DHG-samples) were centrifuged and the supernatants were dialyzed to remove excess glucose, according to the procedures described above. After dialysis, the pellets and supernatants were again combined before being freeze dried using an Edwards Lyofast S 08 Freeze Dryer (A. de Jong TH, Papendrecht, The Netherlands). All freeze-dried samples were milled to fine powder using a knife-type laboratory mill (IKA A11, Staufen, Germany) before further analysis.

**Protein solubility, protein content and LPS contamination**

Solubility of each of the protein samples was assessed creating 1 mg/mL solutions in DPBS (Sigma Aldrich; D8537-500mL). After 30 minutes, samples were centrifuged at 300 × g for 5 min at room temperature and supernatants were transferred to a new tube. Tubes were weighted before and after centrifugation and transfer of the supernatant, and the presence or absence of a pellet was calculated based on weight balance. Supernatant was subsequently checked for Lipopolysaccharide (LPS) contamination using the Pierce LAL Chromogenic Endotoxin Quantitation Kit, according to the manufacturer's instructions (**Table S2**), and proteins in both supernatant and pellets were visualized using SDS PAGE electrophoresis. Protein content of peanut extracts (P-D/DH/DHG) were measured using Pierce™ BCA Protein Assay Kit (Thermo Scientific, 23227).

**SDS PAGE electrophoresis and solubility analysis**

Soluble and insoluble protein fractions resulting from the solubility test were loaded on Biorad Any kD Mini-PROTEAN TGX precast gels (4569035) using reducing (with 10% added β-mercapto-ethanol) or non-reducing conditions. Around 10 µg of soluble protein was mixed 1:1 with 2x Laemmli sample buffer and boiled for 5 min. before loading. Pellets were boiled 30 min. before loading to increase solubility. Precision Plus Protein Dual Xtra Prestained Protein Standard was used as molecular weight (Mw) marker (161-0377; Bio-Rad Laboratories B.V., The Netherlands). Gels were stained overnight using 0.025% Coomassie staining solution and subsequently destained in a methanol/acetic acid solution (MeOH:Hac:H20 in a ratio of 1:1:8). Gels were photographed and analyzed using the Universal Hood III gel doc imaging system (Bio-Rad Laboratories B.V., The Netherlands). Based on size distribution, the peanut extract contained all relevant major peanut allergens (Ara h 1, Ara h 2, Ara h 3 and Ara h 6 (**Figure S1**).

**Detailed Experimental Section 2: Autologous moDC-T cell and B cell assay**

**PBMC isolation**

Peripheral blood mononuclear cells (PBMCs) were isolated from ~70 mL of heparinized whole blood, diluted 1:1 with wash buffer (PBS + 2% Fetal Bovine Serum (FBS) (Biowest, S1810-

1-500U, S00Q3), and layered into Leucosep tubes (Greiner, 227288). After centrifugation (13 min, 1000 × g, 20 °C, acceleration/deceleration 4), the PBMC layer was collected and washed twice with wash buffer (10 min, 612 × g and 300 × g, respectively, at 20 °C).

**Monocyte, T cell and B cell isolation**

Directly after PBMC isolation, CD14⁺ monocytes (positive selection), CD19⁺ B cells (positive selection), and CD4⁺ T cells (negative selection) were sequentially isolated from the same PBMC sample using MACS (Miltenyi Biotec kits: 130-050-201, 130-050-301, 130-096-533) following the manufacturer’s protocol with isolation buffer (PBS + 0.5% BSA + 2 mM EDTA). Cells were stored in liquid nitrogen until further use. Monocytes and moDCs were cultured in RPMI + 10% FBS + 1% pen/strep (= moDC culture medium). CD4⁺ T cells and CD19⁺ B cells were cultured in IMDM + 5% FBS, 1% pen/strep, 20 µg/mL apo-transferrin (Sigma–Aldrich, T1147-100MG), and 50 µM *β*-mercaptoethanol (Gibco, 31350-010) (= T cell culture medium).

**MoDC maturation and differentiation**

CD14⁺ monocytes were differentiated into moDCs over six days in RPMI 1640 (Lonza) supplemented with 10% FBS and 1% pen/strep, using GM-CSF (60 ng/mL; Prospec, CYT221) and IL4 (100 ng/mL; Prospec, CYT211). Medium was refreshed every other day. After six days, immature moDCs were centrifuged (300 × g, 10 min, RT), and one-third of the supernatant was replaced with fresh medium. Unstimulated controls remained untreated, while all other conditions received a type 2 polarizing DC2 mix (1 μg/mL prostaglandin E2 (Prospec, P0409-1MG), 50 ng/mL tumor necrosis factor-𝛼 (Prospec, CYT-223-a), 25 ng/mL IL1𝛽 (Peptrotech, 200–01b-10ug), 10 ng/mL IL6 (Prospec, CYT-213)) combined with either 10 µg protein/mL of peanut protein (P-D/DH/DHG), or 10 µg protein/mL hypoallergenic protein (Rubisco, Gelatin), or 10 µg powder/mL of four different test samples (**Table 2**). A DC2 plus LPS (*Escherichia coli* O111:B4, Invivogen) control was included to match the highest LPS level (of Rubisco; 60.85 EU/mg). Cells were incubated for 48 h at 37 °C, 5% CO₂.

**Autologous MoDC - T cell co-culture**

After 48 h, maturated moDC were centrifuged (300 × g, 5 min, RT) and supernatants stored at -20 °C for cytokine/chemokine analysis. MoDCs were resuspended in 300 µL cold T cell medium, and 300,000 cells were transferred to a 96-well FACS plate for flow cytometry. The remaining 100,000 moDCs were diluted, and 60,000 were seeded in 600 µL T cell medium in a 12-well suspension plate. Autologous CD4⁺ T cells were thawed and co-cultured at a 1:10 moDC:T cell ratio (600,000 T cells in 600 µL). If fewer cells were available, the numbers and volume were adjusted accordingly for all conditions for this independent donor. T cell-only controls included unstimulated and anti-CD3/CD28-stimulated conditions (data not shown). Co-cultures were incubated for five days at 37 °C, 5% CO₂ without medium change.

**Intracellular T cell staining and autologous T cell supernatant - B cell culture**

After five days, T cells were centrifuged (300 × g, 10 min, RT). Supernatants were stored at -20 °C for cytokine analysis, and 150 µL was transferred to a sterile 96-well U-bottom plate and kept at 37 °C, 5% CO₂ for the B cell assay. Half of the T cells were used for flow cytometry (extracellular and intranuclear staining), while the other half was restimulated with PMA (Sigma–Aldrich, 79346-1MG; 5 ng/mL), and ionomycin (Sigma–Aldrich, I0634-1MG; 750 ng/mL) in presence of GolgiPlug (BD Biosciences, 555029; 1 μL/mL) for 5 h at 37 °C, followed by intracellular cytokine staining (see below). B cells were thawed, washed, and 100,000 cells in 50 µL T cell medium were added to the 150 µL T cell supernatant. All peanut reference or test samples (10 µg/mL) were added to corresponding wells. Controls included B cells stimulated with anti-IgM (Sigma-Aldrich, I0759-1MG; 5 µg/mL), anti-IgM + anti-CD40 (Biolegend, 334302; 50 ng/mL) + IL4 (Prospec, CYT211; 50 ng/mL), anti-IgE (Bio-Rad, STAR147; 10 µg/mL), or left untreated (data not shown). Co-cultures were incubated for 10 days at 37 °C, 5% CO₂, with a medium refresh (with food samples) on day 6. After 10 days, supernatants were stored at -20 °C and B cells were collected for flow cytometry.

For one donor, we did not have enough cells to include all samples in the moDC-T cell co-culture step; therefore, we included only one hypoallergenic condition, rubisco, and not gelatin, for this donor.

**Supplementary Table 1.** Detailed Allergy Explorer (ALEX) multiplex array results of peanut-allergy.

| **Allergy Explorer (ALEX) multiplex array results (kU_A_/L)** | | | | | | |
| --- | --- | --- | --- | --- | --- | --- |
| *Peanut-allergens* | | | | | | |
| *Ara h 1* | *Ara h 2* | *Ara h 3* | *Ara h 6* | *Ara h 8* | *Ara h 9* | *Ara h 15* |
| 7.66 | 9.18 | 19.10 | 12.64 | < 0.1 | < 0.1 | < 0.1 |
| 7.36 | 16.52 | 3.00 | 9.64 | 0.51 | < 0.1 | < 0.1 |
| 7.12 | 7.10 | 0.12 | 10.50 | < 0.1 | < 0.1 | < 0.1 |
| 2.99 | 5.23 | 1.24 | 7.05 | < 0.1 | < 0.1 | < 0.1 |
| 29.63 | 28.79 | 20.45 | 23.33 | < 0.1 | < 0.1 | < 0.1 |
| 48.94 | 48.73 | 39.69 | 46.78 | 0.10 | < 0.1 | < 0.1 |

Each row represents a different donor.

**Supplementary Table 2.** Overview of peanut reference samples and four new dietary proteins that were evaluated; protein-, solubility- and LPS content (in 1 mg/mL solution; 1 EU~ 0.1 ng LPS).

| **Name** | **Protein content (%)** | **Solubility (%)** | **LPS EU/mg**  **protein**  **(mean ± SD)^a^** | **Added protein concentration (µg/mL)**  **Total Soluble** | | **LPS pg/mL per sample added to cells (mean ± SD)^a^** |
| --- | --- | --- | --- | --- | --- | --- |
| *Reference samples* | | | |  |  |  |
| P-D | 87,7 | 96.2 | 1.53 **±** 0.34 | 10 | 9.62 | 1.53 ± 0.34 |
| P-DH | 75,2 | 94.3 | 0.73 **±** 0.19 | 10 | 9.43 | 0.73 ± 0.19 |
| P-DHG | 67,2 | 88.9 | 0.02 **±** 0.008 | 10 | 8.89 | 0.02 ± 0.008 |
| *Test samples* | | | |  |  |  |
| P1 | 10.1 | 9.2 | 283.50 **±** 5.89 | 1.01 | 0.09 | 28.6 ± 0.60 |
| P3 | 46.5 | 28.6 | 0.95 **±** 0.92 | 4.65 | 1.33 | 0.44 ± 0.43 |
| P5 | 75.8 | 46.4 | 5.41 **±** 0.07 | 7.58 | 3.52 | 4.10 ± 0.056 |
| P7 | 80.6 | 47.7 | 33.55 **±** 0.45 | 8.06 | 3.84 | 27.0 ± 0.36 |
| *Hypoallergenic samples* | | | |  |  |  |
| Rubisco | 95.7 | 100 | 60.85 **±** 0.58 | 10 | 10 | 60.85 ± 0.58 |
| Gelatin | 26.9 | 100 | 0.12 **±** 0.005 | 10 | 10 | 0.12 ± 0.005 |

^a^For P-D, P-DH, P-DHG and hypoallergenic proteins, samples were added based on BCA-determined protein concentration (10 µg protein/mL), whereas for P1, P3, P5 and P7, samples were added based on powder weight (10 µg powder/mL), with the actual protein content of the added powder determined by Dumas analysis. The soluble plus insoluble part was added. Reported LPS values (EU/mg) represent endotoxin content normalized per mg protein. The LPS concentration (pg/mL) added to the cells was calculated directly from the protein concentration used (P-D, P-DH, P-DHG and hypoallergenic proteins) or corrected for protein content (%, Dumas) of the powder added (P1, P3, P5 and P7).

**Supplementary Table 3.** PBMC panel for flow cytometry to characterize PBMC composition**.**

| **Name** | **Target** | **Brand** | **Cat. nr.** | **Dilution** |
| --- | --- | --- | --- | --- |
| Fixable Viability dye eFluor™ 780 (FVD) | Viability | Thermo Fisher Scientific | 65-0865 | 2000x |
| CD4 monoclonal antibody (OKT4) PerCP-Cyanine5.5 | CD4 | Thermo Fisher Scientific | 45-0048 | 80x |
| PE anti-human CD45RA Antibody (HI100) | CD45RA | BioLegend | 304107 | 320 |
| Brilliant Violet 421™ anti-human CD45RO Antibody (UCHL1) | CD45RO | BioLegend | 304223 | 320 |
| Brilliant Violet 510™ anti-human CD197 (CCR7) Antibody (G043H7) | CD197 (CCR7) | BioLegend | 353232 | 20 |

**Supplementary Table 4.** Corresponding isotype antibodies for the PBMC panel for flow cytometry to characterize PBMCs**.**

| **Isotype** | **Brand** | **Cat. Nr.** |
| --- | --- | --- |
| Mouse IgG2b kappa Isotype Control (eBMG2b), PerCP-Cyanine5.5, eBioscience™ | Thermo Fisher Scientific | 45-4732 |
| PE Mouse IgG2b, κ Isotype Ctrl Antibody (MPC-11) | BioLegend | 400313 |
| Brilliant Violet 421™ Mouse IgG2a, κ Isotype Ctrl Antibody (MOPC-173) | BioLegend | 400259 |
| Brilliant Violet 510™ Mouse IgG2a, κ Isotype Ctrl Antibody (MOPC-173) | BioLegend | 400267 |

**Supplementary Table 5.** MoDC panel for flow cytometry.

| **Name** | **Target** | **Brand** | **Cat. nr.** | **Dilution** |
| --- | --- | --- | --- | --- |
| FVD | Viability | Invitrogen™ | L34982 | 1100x |
| CD14 Monoclonal Antibody (61D3), PerCP-Cyanine5.5 | CD14 | eBioscience™ Invitrogen | 45-0149-42 | 80x |
| V450 Mouse Anti-Human HLA-DR | HLA-DR | BD Biosciences | 561359 | 80x |
| APC Mouse Anti-Human CD209 | CD209 | BD Biosciences | 551545 | 80x |
| CD80 (B7-1) Monoclonal Antibody (2D10.4) FITC | CD80 | eBioscience™ Invitrogen | 11-0809-42 | 80x |
| CD86 (B7-2) Monoclonal Antibody (IT2.2), PE-Cyanine7 | CD86 | eBioscience™ Invitrogen | 25-0869-42 | 1280x |
| PE Mouse Anti-Human OX40 Ligand (CD252) | OX40L | BD Biosciences | 558164 | 80x |
| Brilliant Violet 510™ anti-human CD197 (CCR7) Antibody (G043H7) | CCR7 | BioLegend | 353232 | 20x |
| BD OptiBuild™ BUV661 Mouse Anti-Human CD274 Clone MIH1 (RUO) | PD-L1 (CD274) | BD Biosciences | 741666 | 80x |
| APC/Fire™ 750 anti-human TSLPR (TSLP-R) Antibody clone 1B4 | TSLP-R (CRLF2) | Biolegend | 322814 | 200x |
| Human ST2/IL-33R Alexa Fluor® 700-conjugated Antibody Clone # 2154E | ST2/IL-33R | R&D systems | FAB10118N-100UG | 120x |
| PE/Dazzle™ 594 anti-human CD184 (CXCR4) Antibody clone 12G5 | CD184 (CXCR4) | Biolegend | 306526 | 80x |

**Supplementary Table 6.** Corresponding isotypes for the moDC panel for flow cytometry.

| **Isotype** | **Brand** | **Cat. Nr.** |
| --- | --- | --- |
| Mouse IgG1 kappa Isotype Control (P3.6.2.8.1), PerCP-Cyanine5.5 | eBioscience™ Invitrogen | 45-4714-82 |
| V450 Mouse IgG2a, κ Isotype Control | BD Biosciences | 560550 |
| APC Mouse IgG2b κ Isotype Control | BD Biosciences | 555745 |
| Mouse IgG1 kappa Isotype Control (P3.6.2.8.1), FITC | eBioscience™ Invitrogen | 11-4714-81 |
| Mouse IgG2b kappa Isotype Control (eBMG2b), PE-Cyanine7 | eBioscience™ Invitrogen | 25-4732-81 |
| PE Mouse IgG1, κ Isotype Control | BD Biosciences | 555749 |
| Brilliant Violet 510™ Mouse IgG2a. κ Isotype Ctrl Antibody | BioLegend | 400267 |
| BD Horizon™ BUV661 Mouse IgG1, κ Isotype Control Clone X40 (RUO) | BD Biosciences | 612966 |
| APC/Fire™ 750 Mouse IgG1, κ Isotype Ctrl Antibody Clone MOPC-21 | Biolegend | 400195 |
| Rat IgG1 Alexa Fluor® 700-conjugated Isotype Control | R&D systems | IC005N |
| PE/Dazzle™ 594 Mouse IgG2a, κ Isotype Ctrl Antibody clone MOPC-173 | Biolegend | 400275 |

**Supplementary Table 7.** Extracellular and intranuclear T cell staining for flow cytometry, without prior restimulation of the T cells.

| **Name** | **Target** | **Brand** | **Cat. nr.** | **Dilution** |
| --- | --- | --- | --- | --- |
| FVD |  | Invitrogen™ | L34982 | 1100x |
| CD4 Monoclonal Antibody (OKT4 (OKT-4)), PerCP-Cyanine5.5, eBioscience™ | CD4 | Thermo Fisher | 45-0048 | 320x |
| BD Pharmingen™ Alexa Fluor® 488 Mouse Anti-Human CD183 (Clone 1C6/CXCR3) | CD183 CXCR3) | BD Biosciences | 558047 | 80x |
| BD Pharmingen™ Alexa Fluor® 647 Rat Anti-Human CD294 | CD294 (CRTH2) | BD Biosciences | 558042 | 100x |
| CD69 Monoclonal Antibody (FN50), PE, eBioscience™ clone  FN50 | CD69 | Invitrogen | 12-0699-42 | 120x |
| CD25 Monoclonal Antibody (BC96), eFluor™ 450eBioscience™cloneBC96 | CD25 | Invitrogen | 48-0259-42 | 100x |
| FOXP3 Monoclonal Antibody (236A/E7), PE-Cyanine7, eBioscience™ | FOXP3 | Invitrogen | 25-4777-42 | 250x |
| Brilliant Violet 510™ anti-human CD154 Antibody clone 24-31 | CD40L (CD154) | Biolegend | 310830 | 100x |
| APC/Cyanine7 anti-human CD185 (CXCR5) Antibody clone J252D4 | CXCR5 (CD185) | Biolegend | 356926 | 250x |
| BD Horizon™ BUV661 Mouse Anti-Human CD196 (CCR6) Clone 11A9 (RUO) | CD196 (CCR6) | BD Biosciences | 569509 | 100x |
| PE/Dazzle™ 594 anti-human CD194 (CCR4) Antibody clone L291H4 | CCR4 (CD194) | Biolegend | 359420 | 200x |
| BD Horizon™ BV650 Mouse Anti-Human CD161 Clone DX12 (RUO) | CD161 | BD Biosciences | 563864 | 100x |
| BD Horizon™ BUV395 Mouse Anti-Human CD45RO | CD45RO | BD Biosciences | 564291 | 120x |
| BD Horizon™ BV605 Mouse Anti-Human CD45RA | CD45RA | BD Biosciences | 562886 | 150x |

**Supplementary Table 8.** Corresponding isotype antibodies for extracellular and intranuclear T cell staining for flow cytometry, without prior restimulation of the T cells.

| **Isotype** | **Brand** | **Cat. Nr.** |
| --- | --- | --- |
| Mouse IgG2b kappa Isotype Control (eBMG2b), PerCP-Cyanine5.5, eBioscience™ | Thermo Fisher Scientific | 45-4732-82 |
| Mouse IgG1 kappa Isotype Control (P3.6.2.8.1), Alexa Fluor™ 488, eBioscience™ | Thermo Fisher Scientific | 53-4714 |
| BD Pharmingen™ Alexa Fluor® 647 Rat IgG2a, κ Isotype Control Clone R35-95 (RUO) | BD Biosciences | 557690 |
| Mouse IgG1 kappa Isotype Control (P3.6.2.8.1), PE, eBioscience™ | Invitrogen | 12-4714-81 |
| Mouse IgG1 kappa Isotype Control (P3.6.2.8.1), eFluor™ 450, eBioscience™ | Invitrogen | 48-4714-80 |
| Mouse IgG1 kappa Isotype Control (P3.6.2.8.1), PE-Cyanine7, eBioscience™ | Invitrogen | 25-4714-80 |
| Brilliant Violet 510™ Mouse IgG1, κ Isotype Ctrl Antibody clone MOPC-21 | Invitrogen | 400171 |
| APC/Cyanine7 Mouse IgG1, κ Isotype Ctrl Antibody clone MOPC-21 | Invitrogen | 400127 |
| BD Horizon™ BUV661 Mouse IgG1, κ Isotype Control Clone X40 (RUO) | BD Biosciences | 612966 |
| PE/Dazzle™ 594 Mouse IgG1, κ Isotype Ctrl Antibody clone MOPC-21 | Biolegend | 400175 |
| BD Horizon™ BV650 Mouse IgG1, k Isotype Control Clone X40 (RUO) | BD Biosciences | 563231 |
| BD Horizon™ BUV395 Mouse IgG2a, κ Isotype Control Clone G155-178 (RUO) | BD Biosciences | 563809 |
| BD Horizon™ BV605 Mouse IgG2b, κ Isotype Control | BD Biosciences | 563099 |

**Supplementary Table 9.** Extracellular and intracellular T cell staining for flow cytometry. T cells first need to be restimulated using a PMA, Ionomycin and Golgiplug mixture for 5 hours before measurements.

| **Name** | **Target** | **Brand** | **Cat. nr.** | **Dilution** |
| --- | --- | --- | --- | --- |
| Fixable Viability Dye eFluor™ 780 (APC-Cy7) |  | Thermofisher | 65-0865-14 | 2000x |
| CD4 Monoclonal Antibody (OKT4 (OKT-4)), PerCP-Cyanine5.5, eBioscience™ | CD4 | Thermo Fisher | 45-0048 | 80x |
| Brilliant Violet 421™ anti-human IL10 Antibody | IL10 | BioLegend | 501422 | 160x |
| PE/Cyanine7 anti-human IFN-γ Antibody | IFNy | BioLegend | 502528 | 20x |
| IL13 Monoclonal Antibody (85BRD), PE, eBioscience™ | IL13 | eBioscience | 12-7136-42 | 80x |
| Brilliant Violet 510™ anti-human IL4 Antibody | IL4 | BioLegend | 500836 | 20x |

**Supplementary Table 10.** Corresponding isotype antibodies for extracellular and intracellular T cell panel for flow cytometry. T cells first need to be restimulated using a PMA, Ionomycin and Golgiplug mixture for 5 hours before measurements.

| **Isotype** | **Brand** | **Cat. Nr.** |
| --- | --- | --- |
| Mouse IgG2b kappa Isotype Control (eBMG2b), PerCP-Cyanine5.5, eBioscience™ | Thermo Fisher Scientific | 45-4732-82 |
| Brilliant Violet 421™ Rat IgG1, κ Isotype Ctrl Antibody | BioLegend | 400429 |
| PE/Cyanine7 Mouse IgG1. κ Isotype Ctrl Antibody | BioLegend | 400125 |
| Mouse IgG2b kappa Isotype Control (eBMG2b), PE, eBioscience™ | Thermo Fisher Scientific | 12-4732-81 |
| Brilliant Violet 510™ Rat IgG1. κ Isotype Ctrl Antibody | BioLegend | 400435 |

**Supplementary Table 11.** B cell panel for flow cytometry.

| **Name** | **Target** | **Brand** | **Cat. nr.** | **Dilution** |
| --- | --- | --- | --- | --- |
| FVD |  | Invitrogen™ | L34982 | 1100x |
| FITC anti-human CD19 Antibody HIB19 | CD19 | Biolegend | 302206 | 80x |
| BD Pharmingen™ Alexa Fluor® 700 Mouse Anti-Human CD20 Clone 2H7 (RUO) | CD20 | BD Biosciences | 560631 | 120x |
| CD4 Monoclonal Antibody (RPA-T4), eFluor™ 506, eBioscience™ RPA-T4 | CD4 | Invitrogen | 69-0049-42 | 200x |
| BD Horizon™ BV421 Mouse Anti-Human CD38 Clone HIT2 (RUO) | CD38 | BD Biosciences | 562444 | 100x |
| BD Pharmingen™ APC Mouse Anti-Human CD27 Clone M-T271 (RUO) | CD27 | BD Biosciences | 561400 | 100x |
| CD138 (Syndecan-1) Monoclonal Antibody (DL-101), PE, eBioscience™ clone DL-101 | CD138 | Invitrogen | 12-1389-42 | 100x |
| APC/Cyanine7 anti-human IgG Fc Antibody clone M1310G05 | IgG Fc | Biolegend | 410732 | 250x |
| BD OptiBuild™ BUV661 Mouse Anti-Human IgE clone Clone G7-26 (RUO) | IgE | BD Biosciences | 750573 | 250x |

**Supplementary Table 12.** Corresponding isotype antibodies for B cell panel for flow cytometry.

| **Isotype** | **Brand** | **Cat. Nr.** |
| --- | --- | --- |
| FITC Mouse IgG1, κ Isotype Ctrl Antibody clone MOPC-21 | Biolegend | 400107 |
| BD Pharmingen™ Alexa Fluor® 700 Mouse IgG2b, κ Isotype Control Clone 27-35 (RUO) | BD Biosciences | 560543 |
| Mouse IgG1 kappa Isotype Control (P3.6.2.8.1), eFluor™ 506, eBioscience™ | Invitrogen | 69-4714-80 |
| BD Horizon™ BV421 Mouse IgG1, k Isotype Control Clone X40 (RUO) | BD Biosciences | 562438 |
| BD Pharmingen™ APC Mouse IgG1, κ Isotype Control Clone MOPC-21 (RUO) | BD Biosciences | 554681 |
| Mouse IgG1 kappa Isotype Control (P3.6.2.8.1), PE, eBioscience™ | Invitrogen | 12-4714-82 |
| APC/Cyanine7 Rat IgG2a, κ Isotype Ctrl Antibody | Biolegend | 400523 |
| BD Horizon™ BUV661 Mouse IgG2a, κ Isotype Control Clone G155-178 (RUO) | BD Biosciences | 612982 |

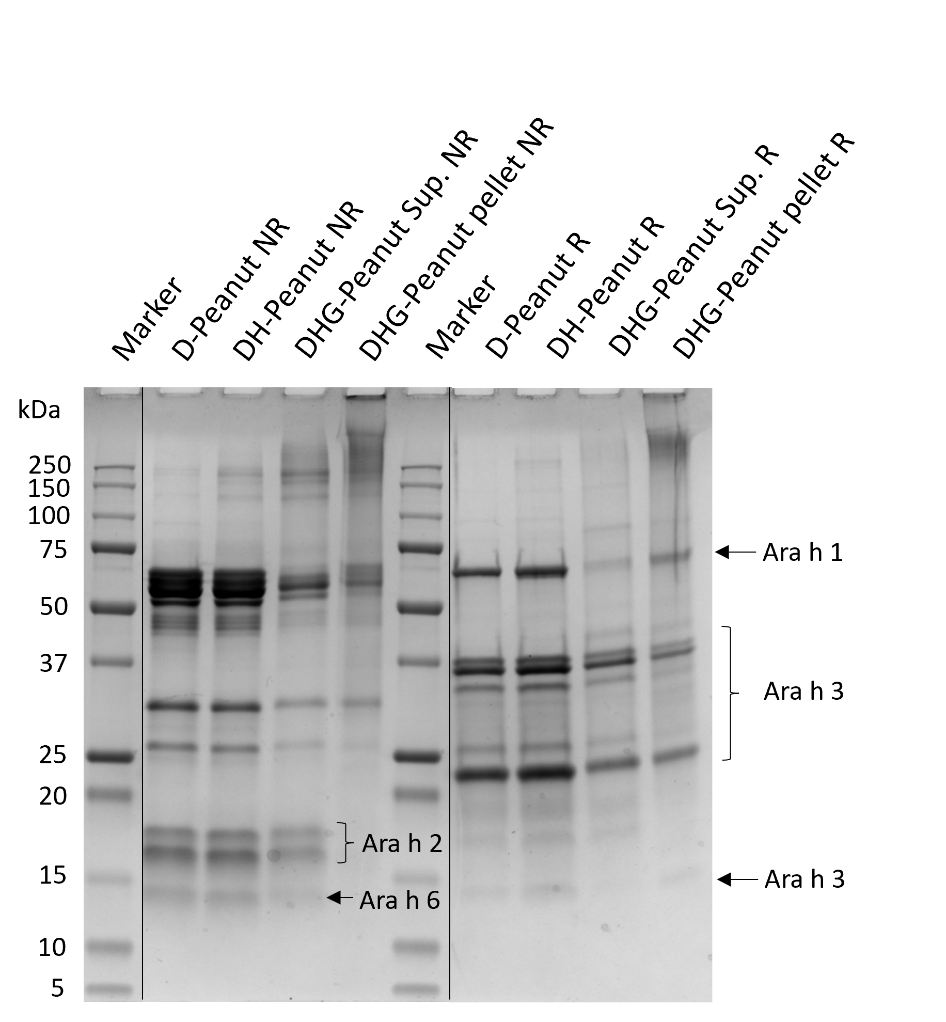

**Supplementary Figure 1.** Peanut protein samples on any kD SDS PAGE using non-reducing (NR) and reducing (R) conditions. Ara h 1 and Ara h 3 proteins are indicated based on expected Mw sizes in reducing SDS PAGE conditions. Ara h 2 and Ara h 6 are indicated based on expected Mw size in non-reducing conditions. D: Dialyzed protein sample; DH: Dialyzed and heated without glucose; DHG: Dialyzed, heated and glycated in the presence of glucose; Sup.: soluble supernatant. Circa 10 µg of protein was run on gel, based on protein solubility. Heated but non-glycated samples remained soluble. Glycation in the presence of the reducing sugar glucose creates soluble and insoluble aggregates, as visualized by the shift in high molecular weight sized bands and smearing on gel. In addition, some large aggregates unable to move into the SDS PAGE gel during electrophoresis can be observed in the slot of the DHG peanut pellet sample.

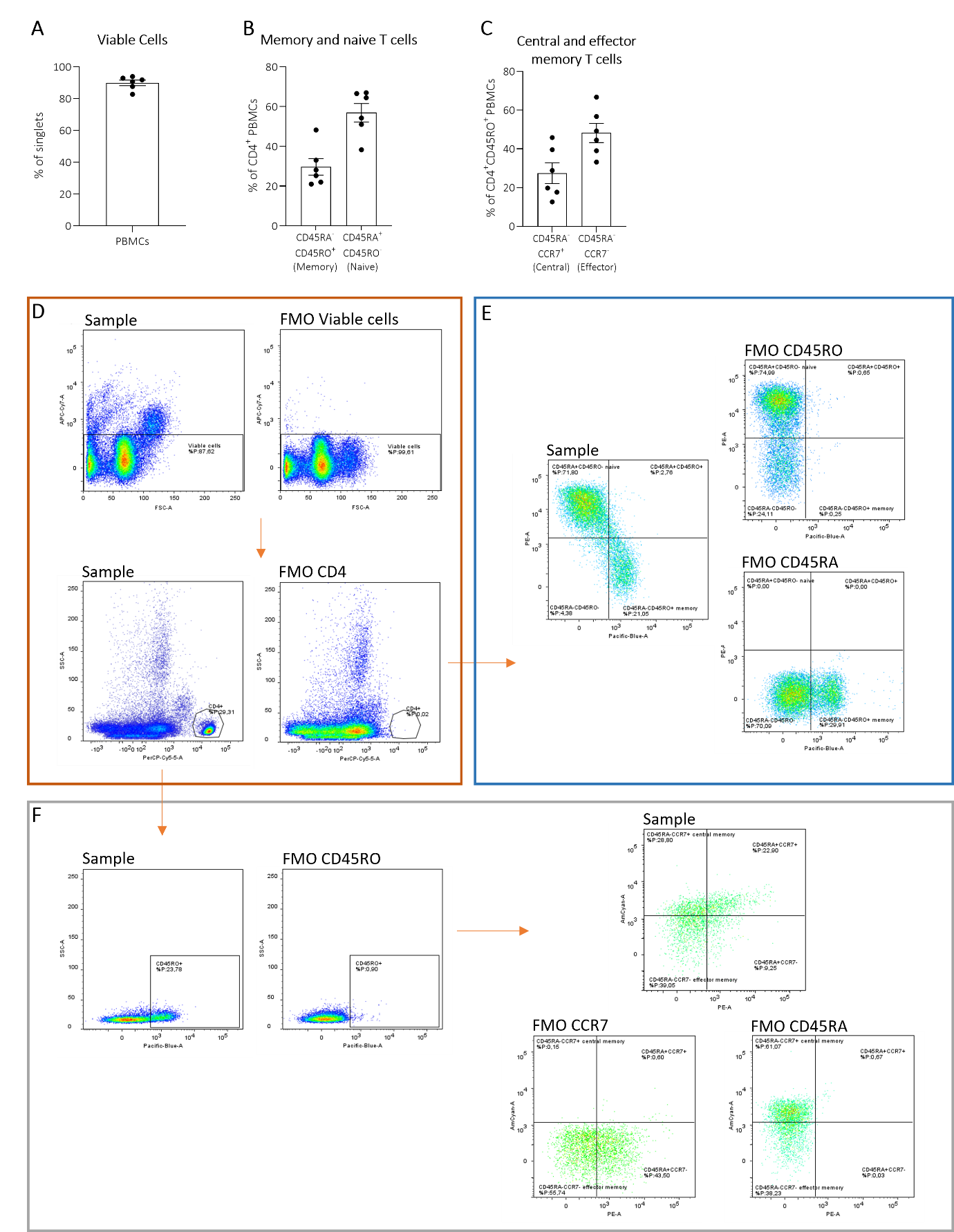

**Supplementary Figure 2. PBMC characterization and flow cytometry gating strategy.** After PBMC isolation, fractions were analyzed for: **A**) viable cells, **B**) memory and naïve CD4⁺ T cells, and **C**) central/effector T cells within CD4⁺CD45RO⁺ cells. **D-F**) Gating strategy of measured PBMC fractions. After sequential gating for constant flow, total cells, and single cells (singlets), the gating strategy was applied as shown above: **D**) within the viable cell population, the CD4⁺ immune cell subset was identified. From CD4⁺ PBMCs, **E**) CD45RO and CD45RA expression, and **F**) CD45RO only expression. CD45RO⁺ cells were then further analyzed for CD45RA and CCR7. FMO= fluorescent minus one control. No statistics were performed as no comparisons were made.

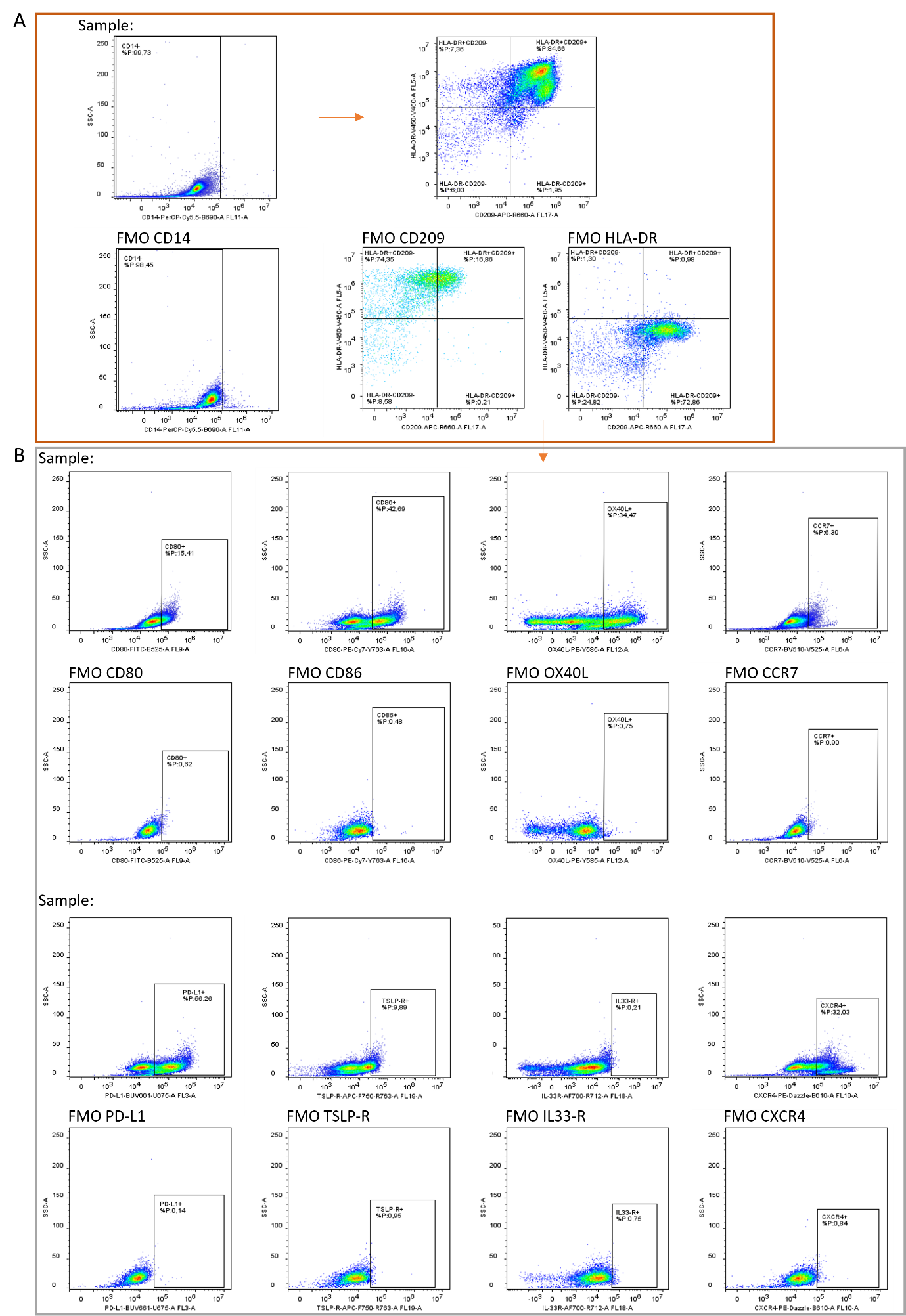

**Supplementary Figure 3.** **MoDC flow cytometry gating strategy.** After 48 hour moDC maturation, moDCs were measured by flow cytometry. After sequential gating for constant flow, total cells, single cells (singlets), and viability, the gating strategy was applied as shown above: **A**) MoDCs were identified as CD14^-^ and CD209^+^HLA-DR^+^. Next, in this CD209^+^HLA-DR^+^ population **B**) CD80, CD86, OX40L, CCR7, PD-L1, TSLP-R, IL33-R, and CXCR4 marker expression were analyzed. FMO= fluorescent minus one control.

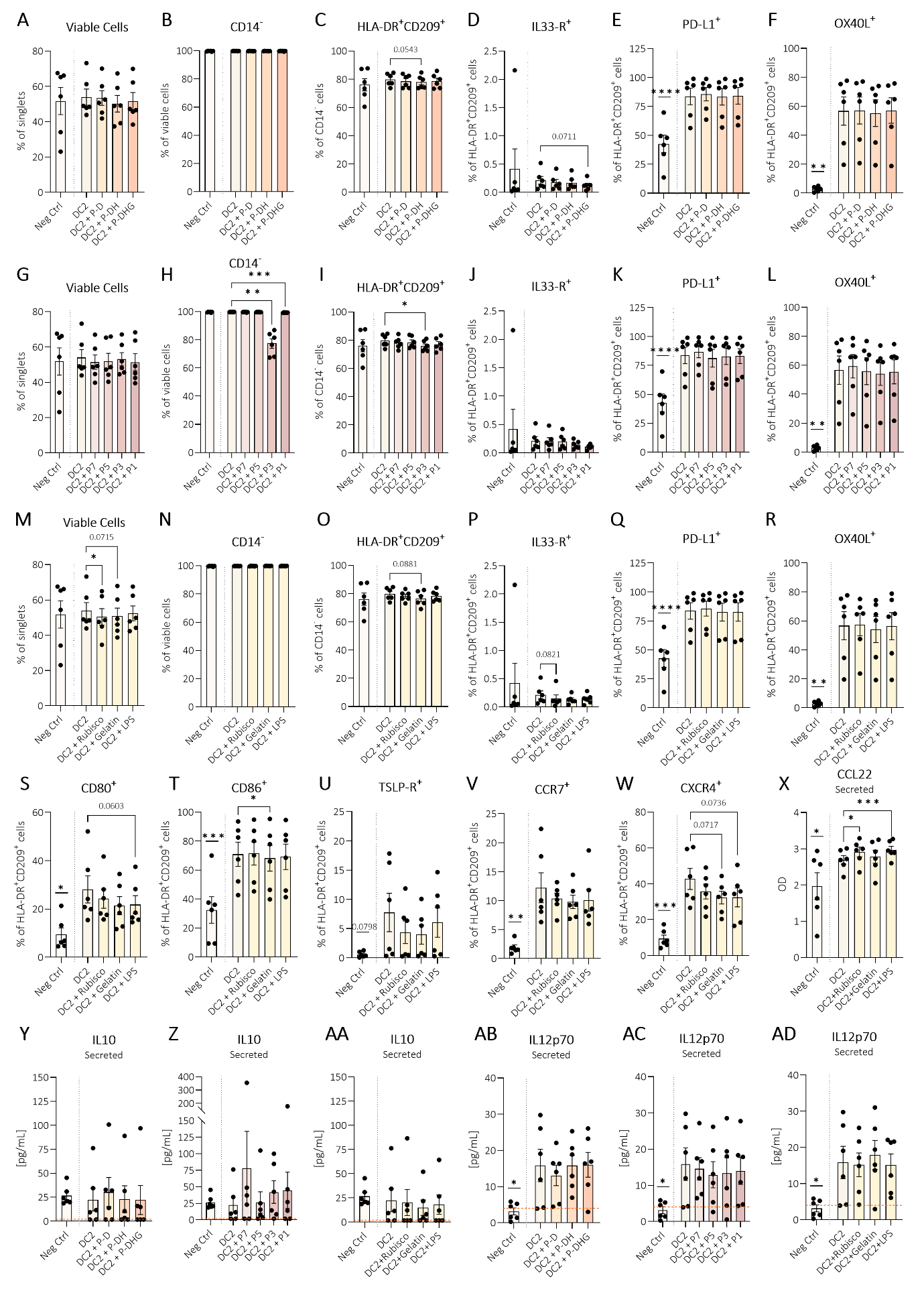
**Supplementary Figure 4.** **MoDC marker expression and cytokine secretion, including controls.** After 48 hours of exposure to peanut (unprocessed/processed) samples, test samples (P7, P5, P3, P1), hypoallergenic samples (rubisco, gelatin), LPS, or left untreated, moDCs were analyzed by flow cytometry and supernatant was analyzed for secreted cytokines. Peanut reference sample group: **A**) Viable cells, **B**) CD14^-^ cells, **C**) CD14^-^HLA-DR^+^CD209^+^ (moDCs), and within the moDC population: **D**) IL33-R, **E**) PD-L1, and **F**) OX40L were measured. Test sample group: **G**) Viable cells, **H**) CD14^-^ cells, **I**) CD14^-^HLA-DR^+^CD209^+^ (moDCs), and within the moDC population: **J**) IL33-R, **K**) PD-L1, and **L**) OX40L were measured. **M-W**) All flow cytometry results for hypoallergenic samples and the LPS control: **M**) Viable cells, **N**) CD14^-^ cells, **O**) CD14^-^HLA-DR^+^CD209^+^ (moDCs), and within the moDC population: **P**) IL33-R, **Q**) PD-L1, **R**) OX40L, **S**) CD80, **T**) CD86, **U**) TSLP-R, **V**) CCR7, and **W**) CXCR4. **X**) Secreted CCL22 controls (OD values). MoDC supernatant was analyzed for secreted IL10 in **Y**) peanut reference samples, **Z**) test samples and **AA**) control stimuli, similarly for IL12p70 in **AB-AD**. Orange dotted lines represent lower detection limit of the ELISA kit. Data were analyzed using paired t-test, Wilcoxon matched-pairs signed rank test, RM one-way ANOVA with Dunnett’s test (with Geisser–Greenhouse correction where applicable), or Friedman test with Dunn’s multiple comparisons. DC2 was compared to unstimulated negative control; Samples (reference, test, hypoallergenic) combined with DC2 were compared to DC2. Bars represent mean ± SEM, N=6, p ≤ 0.05 (*):, p ≤ 0.01 (**), p ≤ 0.001 (***), p ≤ 0.0001 (****).

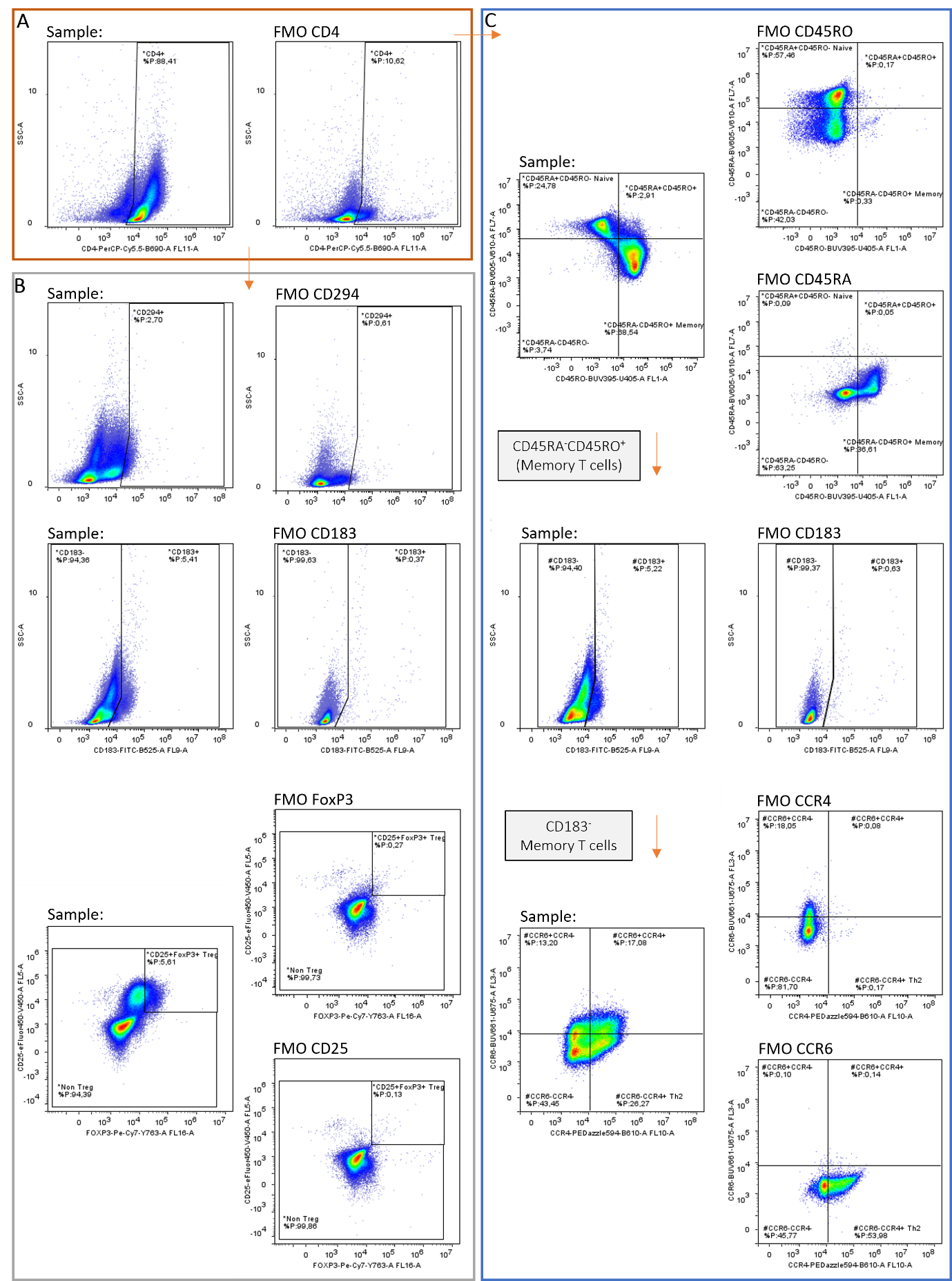

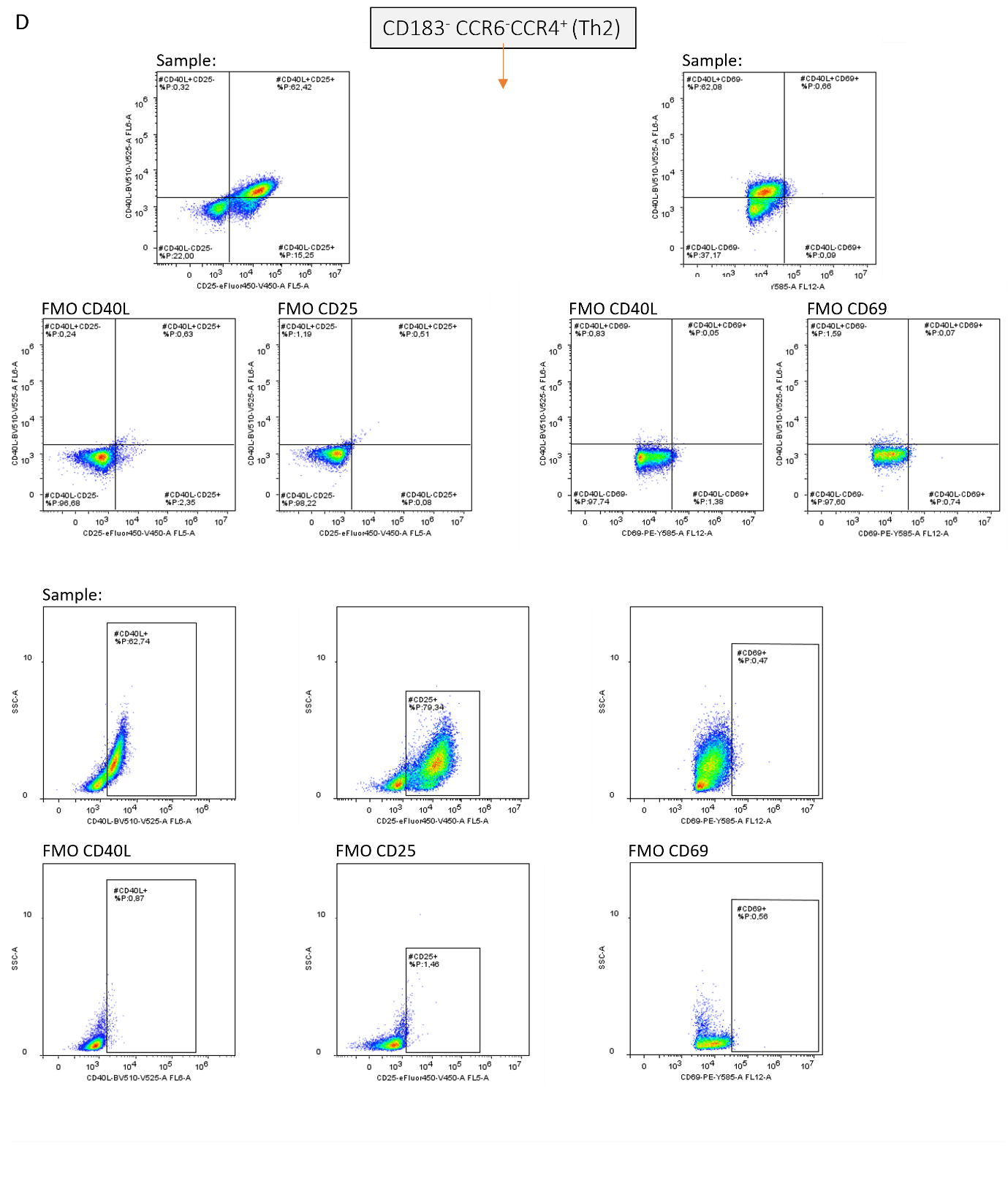

**Supplementary Figure 5.** **Flow cytometry gating strategy to characterize T cell phenotype.** After five days of moDC-T cell co-culture, T cells were measured by flow cytometry. After sequential gating for constant flow, total cells, single cells (singlets), and viability, the gating strategy was applied as shown above: T cells were identified using **A**) CD4 expression. Within this CD4^+^ T cell population, **B**) CD294 (Th2), CD183 (Th1) and CD25^+^FoxP3^+^ (Treg) were identified. **C**) CD45RO and CD45RA were assessed in CD4^+^ cells, followed by gating on memory T cells (CD45RA⁻CD45RO⁺), then CD183⁻ cells, and finally Th2 (CD183⁻CCR6⁻CCR4⁺). In the Th2 population **D**), CD40L, CD25, and CD69 were assessed, including dual expression of CD40L with CD69 or CD25. FMO= fluorescent minus one control.

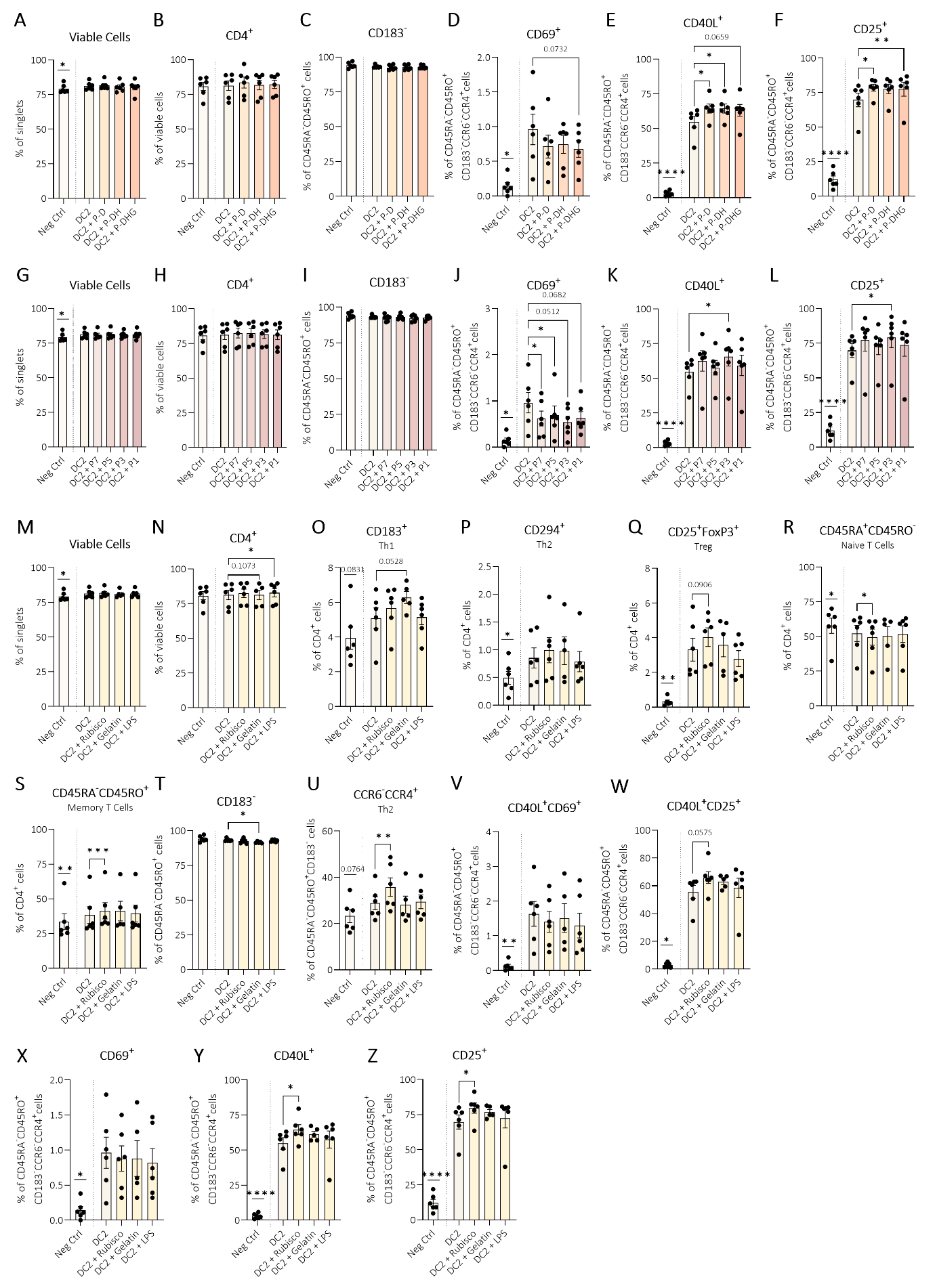
**Supplementary Figure 6**. **T cell phenotype and controls.** After five days of moDC-T cell co-culture, T cells were measured by flow cytometry. Peanut reference sample group: **A**) viable cells, **B**) CD4^+^ T cells, **C**) CD183^-^ T cells (from which Th2 CCR6⁻CCR4⁺ were identified), with expression of **D**) CD69, **E**) CD40L, and **F**) CD25. Test sample group: **G**) viable cells, **H**) CD4^+^ T cells, **I**) CD183^-^ T cells (from which Th2 CCR6⁻CCR4⁺ were identified), with expression of **J**) CD69, **K**) CD40L, and **L**) CD25. **M-Z**) Controls (hypoallergenic, LPS and T cell-only controls) of all measured markers. **M**) Viable cells, **N**) CD4^+^ T cells, and within the CD4^+^ T cell population: **O**) CD183 (Th1), **P**) CD294 (Th2), **Q**) CD25^+^FoxP3^+^ (Treg), **R**) CD45RA^+^CD45RO^-^ (naïve T cells), **S**) CD45RA^-^CD45RO^+^ (memory T cells). From these memory T cells: **T**) CD183^-^, followed by **U**) CD183^-^ CCR6^-^CCR4^+^ (Th2). Within this Th2 population, **V**) CD40L^+^CD69^+^, **W**) CD40L^+^CD25^+^, and **X**) CD69, **Y**) CD40L and **Z**) CD25, were characterized. Data were analyzed using paired t-test, Wilcoxon matched-pairs signed rank test, RM one-way ANOVA with Dunnett’s test (with Geisser–Greenhouse correction where applicable), or Friedman test with Dunn’s multiple comparisons. DC2 was compared to unstimulated negative control; Samples (reference, test, hypoallergenic) combined with DC2 were compared to DC2. Bars represent mean ± SEM, N=6 (except for gelatin; N=5), p ≤ 0.05 (*):, p ≤ 0.01 (**), p ≤ 0.001 (***), p ≤ 0.0001 (****).

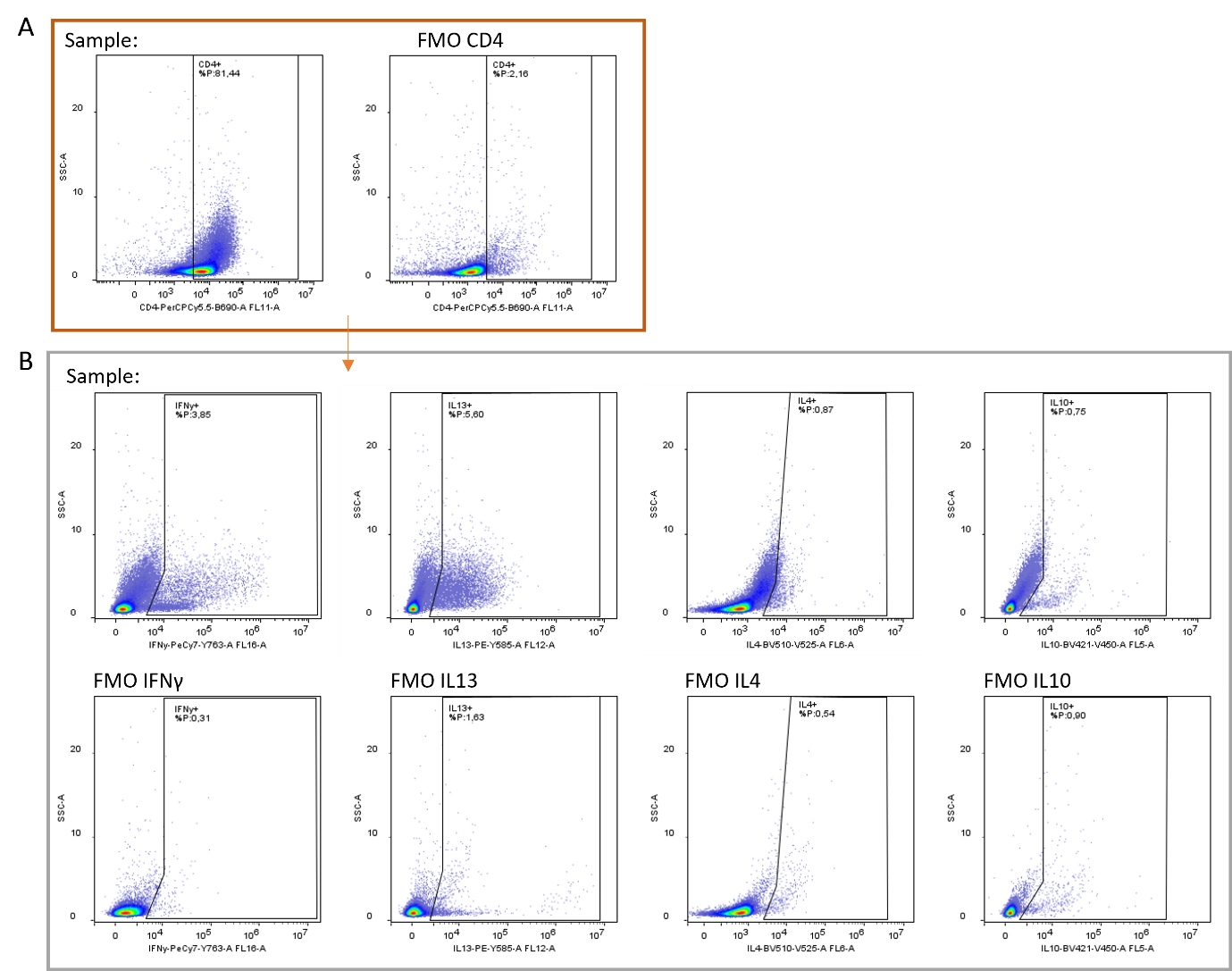
**Supplementary Figure 7. Flow cytometry gating strategy to identify T cell function by intracellular cytokines.** After 5 days of moDC-T cell co-culture, T cells were restimulated for 5 hours before flow cytometry. After sequential gating for constant flow, total cells, single cells (singlets), and viability, the gating strategy was applied as shown above: T cells were identified as **A**) CD4^+^. Next, in this CD4^+^ population **B**) intracellular IFNγ, IL13, IL4, and IL10 expression were analyzed. FMO= fluorescent minus one control.

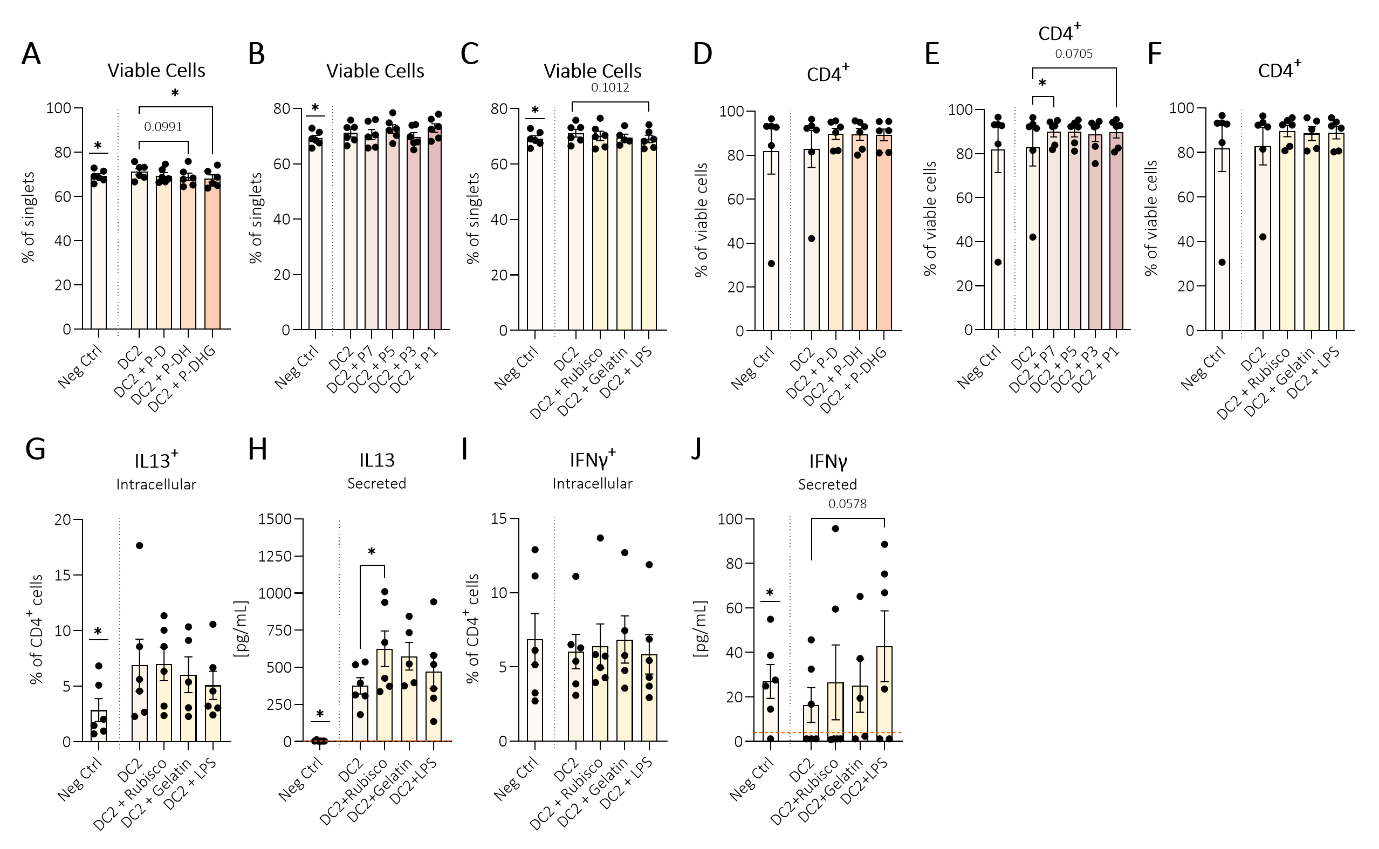
 **Supplementary Figure 8. T cell function characterization (intracellular and secreted cytokines), including controls.** After 5 days of moDC-T cell co-culture, supernatants were collected for ELISA, and T cells were restimulated for 5 hours before flow cytometry. After identifying **A-C**) viable cells, **D-F**) CD4^+^ T cells were characterized within peanut reference samples, test samples and hypoallergenic controls. Controls included hypoallergenic proteins, LPS control and T cell-only controls, in which **G**) intracellular IL13, **H**) secreted IL13, **I**) intracellular IFNγ, and **J**) secreted IFNγ were measured. Data were analyzed using paired t-test, Wilcoxon matched-pairs signed rank test, RM one-way ANOVA with Dunnett’s (with Geisser–Greenhouse correction where applicable), or Friedman test with Dunn’s multiple comparisons. DC2 was compared to unstimulated negative control; Samples (reference, test, hypoallergenic) combined with DC2 were compared to DC2. Bars represent mean ± SEM, N=6 (except for gelatin; N=5), p ≤ 0.05 (*).

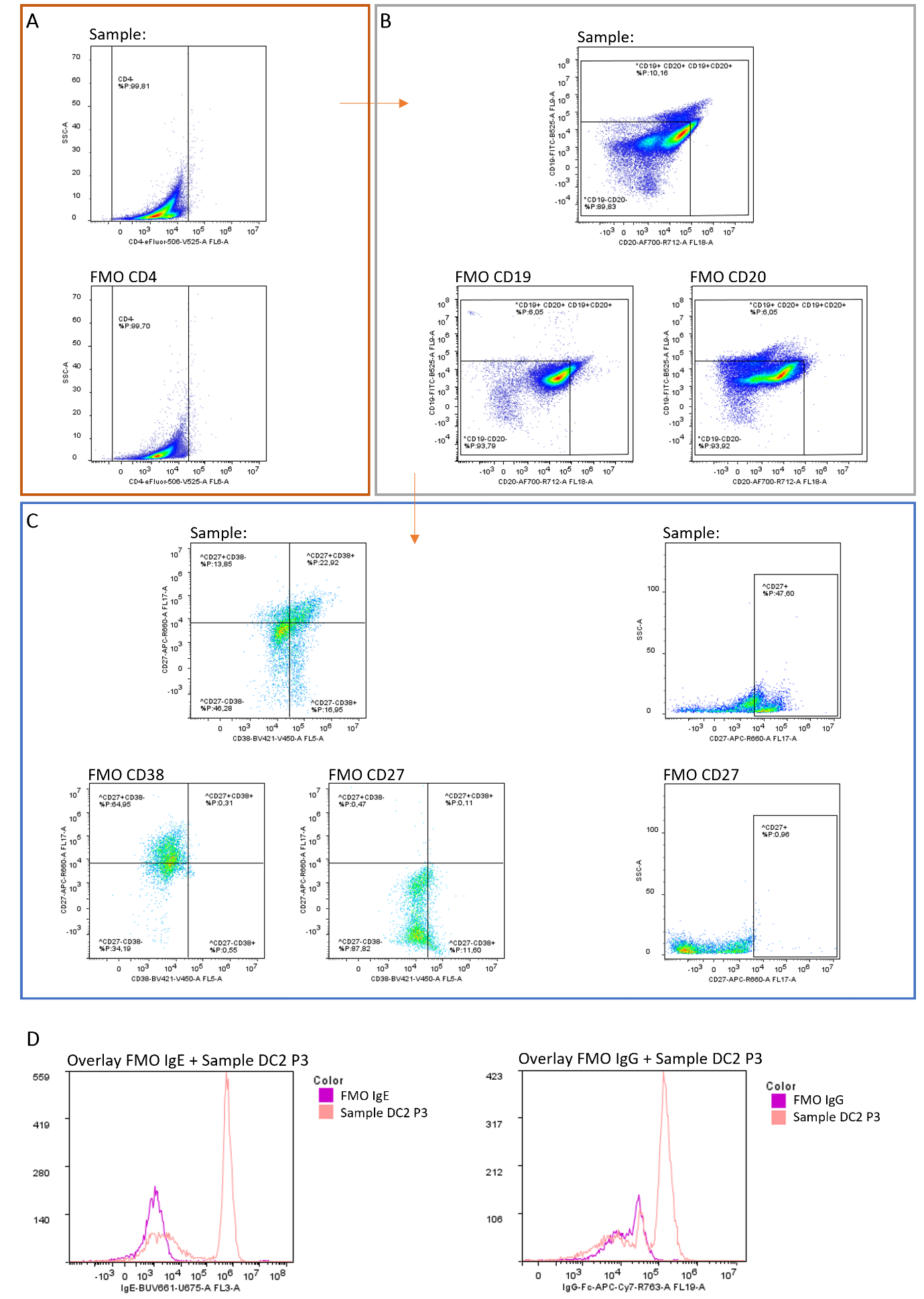

**Supplementary Figure 9. Flow cytometry gating strategy to identify B cell populations.** B cells were cultured in moDC/T cell supernatant in presence of the samples for 10 days and were then measured with flow cytometry. After sequential gating for constant flow, total cells, single cells (singlets), and viability, the gating strategy was applied as shown above: **A**) absence of CD4 was followed by **B**) CD19 and CD20 identification in which all CD19^+^, CD20^+^ and CD19^+^CD20^+^ were gated as one population. Within this CD19/CD20 population, **C**) CD27^+^ and CD27^+^CD38^+^ B cells were identified. As IgE and IgG responder percentages were low (gating/data not shown), MFIs were used; MFIs were also low, therefore only one example of one responding donor/sample is shown; **D**) Example MFI overlays of IgE and IgG for DC2 P3 and the corresponding FMO. FMO= fluorescent minus one control.

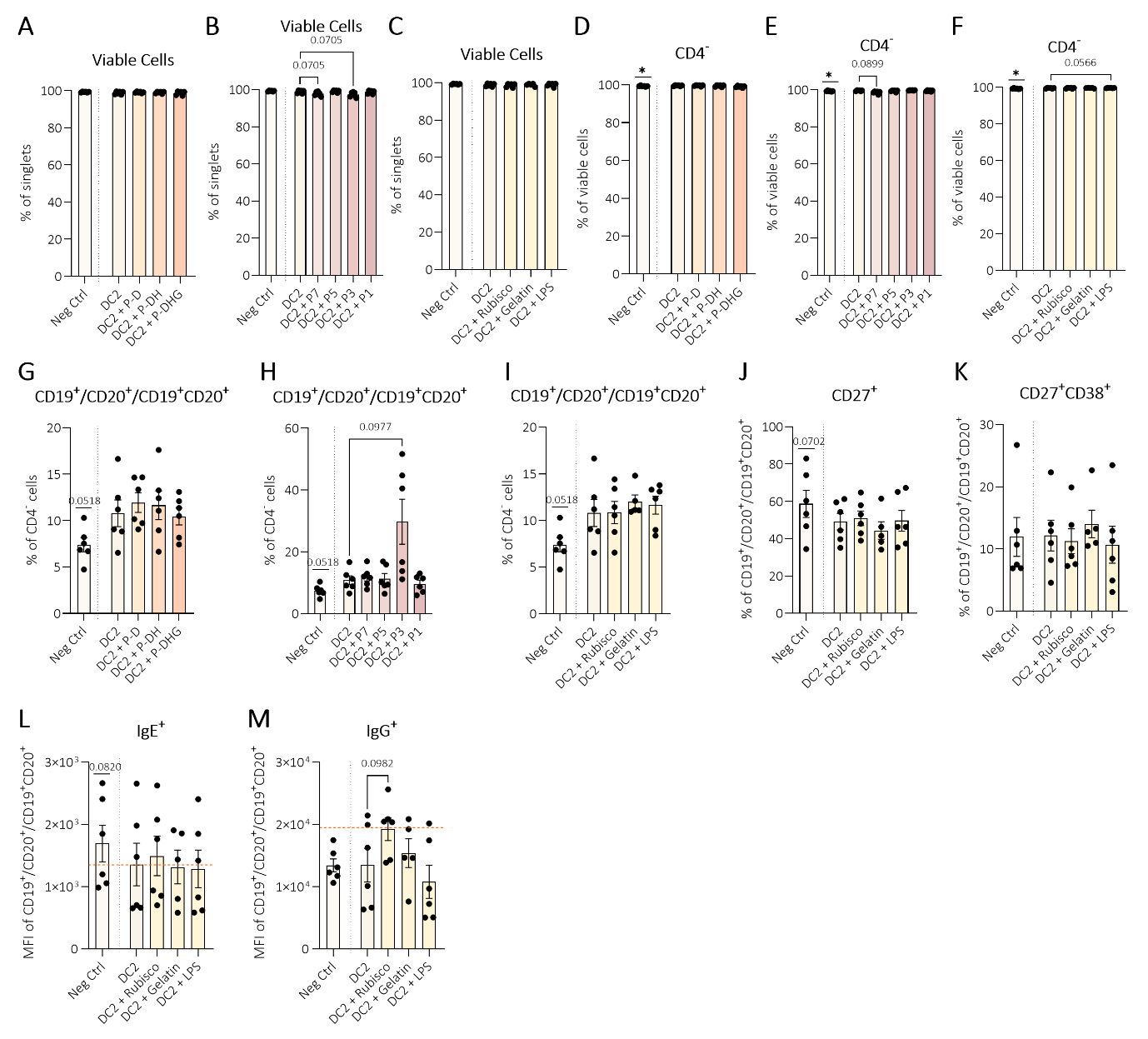

**Supplementary Figure 10.** **B cell characterization and controls.** B cells were cultured in moDC/T cell supernatant in presence of the proteins for 10 days and were then measured with flow cytometry. After **A-C**) viable cells were identified in peanut reference, test, and hypoallergenic groups, **D-F**) CD4⁻ cells were gated. Subsequently, **G-I**) CD19^+^/CD20^+^/CD19^+^CD20^+^ population were characterized and, in this population, **J**) CD27^+^, **K**) CD27^+^CD38^+^, **L**) IgE (MFI), and **M**) IgG (MFI) were identified (in the hypoallergenic group). Orange dotted lines represents average MFI of corresponding FMO. Data were analyzed using paired t-test, RM one-way ANOVA with Dunnett’s test (with Geisser–Greenhouse correction where applicable), or Friedman test with Dunn’s multiple comparisons. DC2 was compared to unstimulated negative control; Samples (reference, test, hypoallergenic) combined with DC2 were compared to DC2. Bars represent mean ± SEM, N=6 (except for gelatin; N=5), p ≤ 0.05 (*), p ≤ 0.01 (**), p ≤ 0.0001 (****).

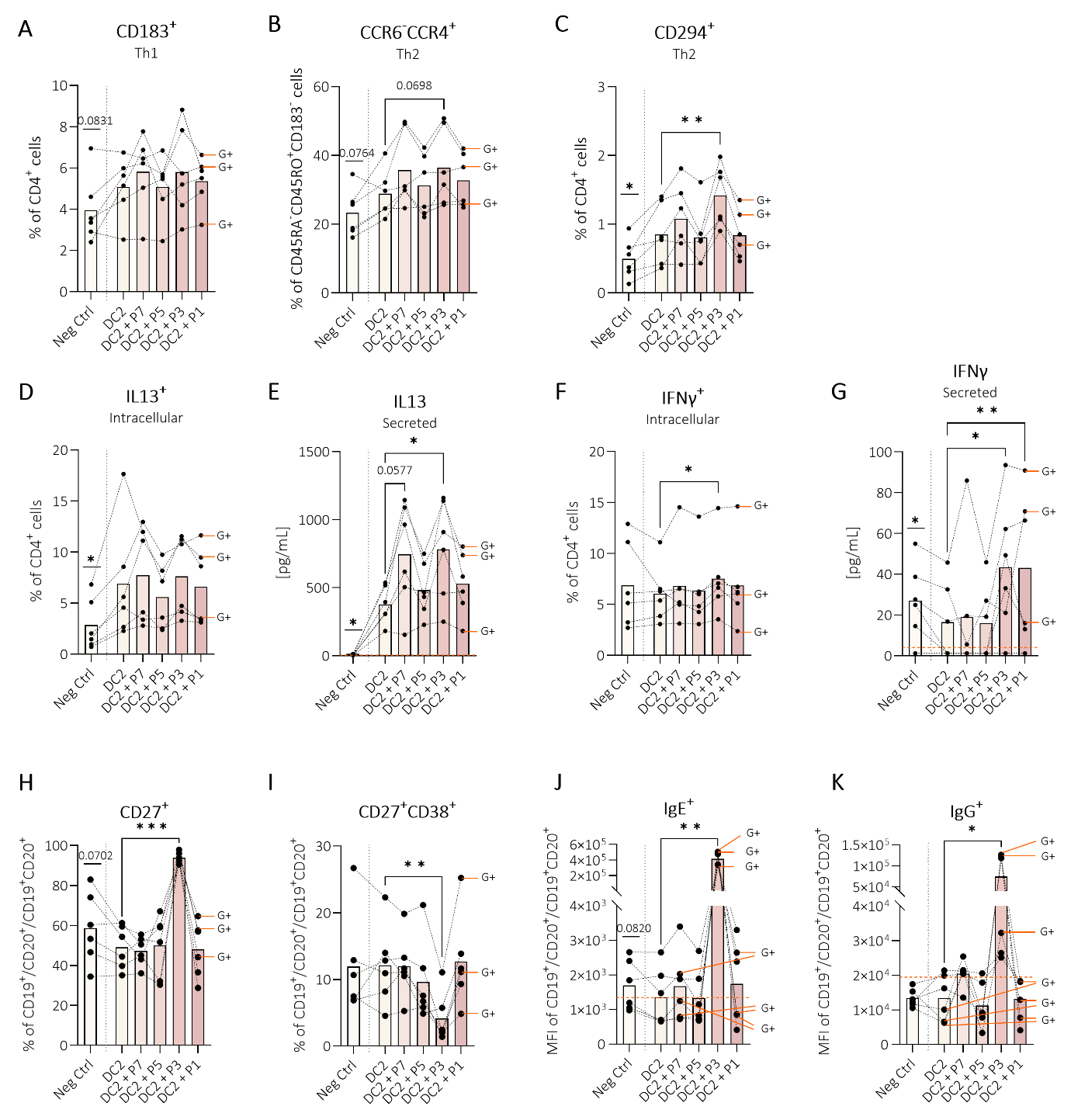

**Supplementary Figure 11. Connected dot plots showing individual donor responses in Th1/Th2 and B cell responses.** The corresponding data are shown in the main manuscript; here, (**A**-**K**) individual donor lines are displayed to provide deeper insight into donor-specific responses to test samples. In addition, donors that were identified as grain-sensitized based on the Allergy Explorer (ALEX) multiplex array were manually labeled (G+) in the graphs above. Orange dotted lines indicate the mean MFI of the corresponding FMO control or ELISA detection limit. DC2 was compared to unstimulated negative control; Test samples combined with DC2 were compared to DC2. Statistical analyses are as shown in the corresponding graphs in the main manuscript. Bars represent the mean; N = 6. p ≤ 0.05 (*), p ≤ 0.01 (**), p ≤ 0.001 (***).

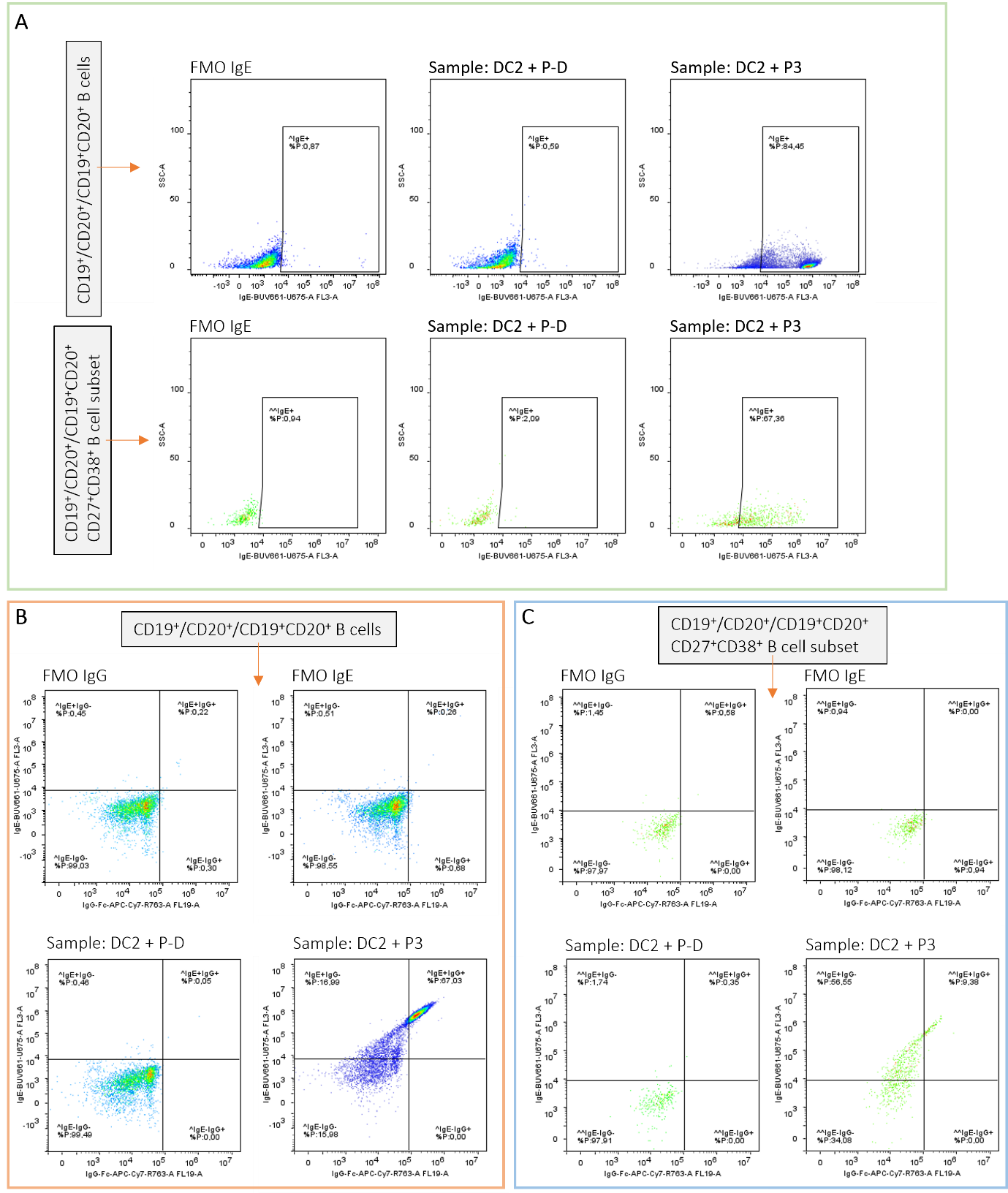

**Supplementary Figure 12. Additional gating strategy to clarify IgE and IgG expression. A**) Gating strategy for intracellular IgE expression in CD19^+^/CD20^+^/CD19^+^CD20^+^ B cells and in the CD27^+^CD38^+^ B cell subset of CD19^+^/CD20^+^/CD19^+^CD20^+^ B cells. **B**) B cells dual positive for IgE and IgG within CD19^+^/CD20^+^/CD19^+^CD20^+^ B cells and **C**) within the CD27^+^CD38^+^ B cell subset of CD19^+^/CD20^+^/CD19^+^CD20^+^ B cells.

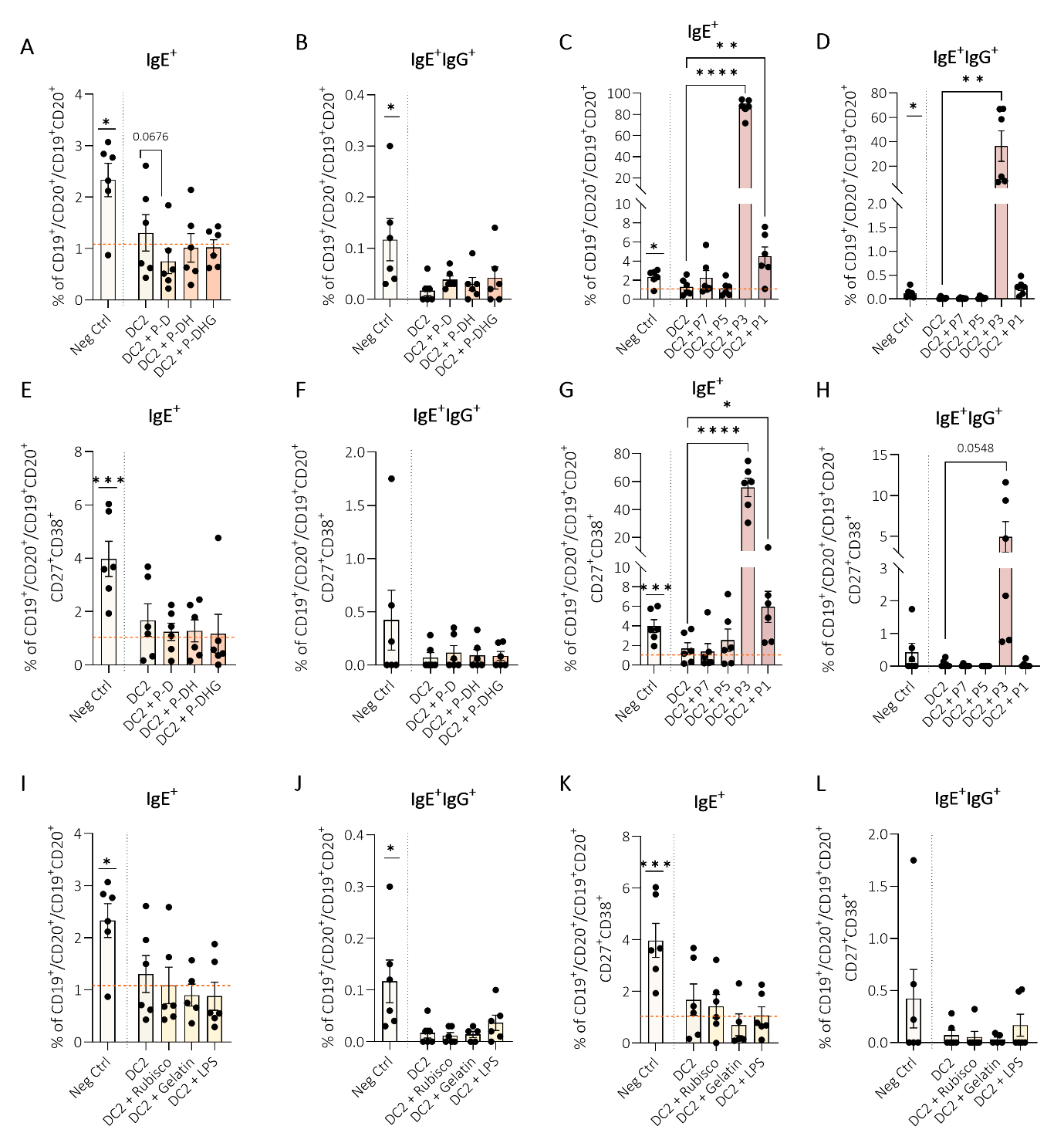

**Supplementary Figure 13. IgE^+^ and IgE^+^IgG^+^ dual positive B cells.** Intracellular percentage of **A**) IgE and **B**) IgE^+^IgG^+^ in CD19^+^/CD20^+^/CD19^+^CD20^+^ B cells exposed to peanut samples, and **C**) IgE and **D**) IgE^+^IgG^+^ in CD19^+^/CD20^+^/CD19^+^CD20^+^ B cells exposed to test samples. Intracellular percentage of **E**) IgE and **F**) IgE^+^IgG^+^ in the CD27^+^CD38^+^ B cell subset exposed to peanut samples, and **G**) IgE and **H**) IgE^+^IgG^+^ in the CD27^+^CD38^+^ B cell subset exposed to test samples. Intracellular **I**) IgE and **J**) IgE^+^IgG^+^ in CD19^+^/CD20^+^/CD19^+^CD20^+^ B cells, and **K**) IgE and **L**) IgE^+^IgG^+^ in the CD27^+^CD38^+^ B cell subset, exposed to hypoallergenic and LPS control samples. As seen in the gating strategy of the CD27^+^CD38^+^ B cell subset in **Figure S11**, results should be interpreted with caution due to the low number of cell events in this B cell subset. Orange dotted lines represents average % of corresponding FMO. Data were analyzed using paired t-test, Wilcoxon matched-pairs signed rank test, RM one-way ANOVA with Dunnett’s test, mixed-effects models with Dunnett’s test, or Friedman test with Dunn’s multiple comparisons. DC2 was compared to unstimulated negative control; Samples (reference, test, hypoallergenic) combined with DC2 were compared to DC2. Bars represent mean ± SEM, N=6 (except for gelatin; N=5), p ≤ 0.05 (*), p ≤ 0.01 (**), p ≤ 0.001 (***), p ≤ 0.0001 (****).
